# Diverse routes of Club cell evolution in lung adenocarcinoma

**DOI:** 10.1101/2021.06.10.447936

**Authors:** Yuanyuan Chen, Reka Toth, Sara Chocarro, Dieter Weichenhan, Joschka Hey, Pavlo Lutsik, Stefan Sawall, Georgios T. Stathopoulos, Christoph Plass, Rocio Sotillo

## Abstract

The high plasticity of lung epithelial cells, has for many years, confounded the correct identification of the cell-of-origin of lung adenocarcinoma (LUAD), one of the deadliest malignancies worldwide. Here, we address the cell-of-origin of LUAD, by employing lineage-tracing mouse models combined with a CRISPR/Cas9 system to induce an oncogenic *Eml4-Alk* rearrangement in virtually all epithelial cell types of the lung. We find that Club cells give rise to lung tumours with a higher frequency than AT2 cells. Based on whole genome methylome, we identified that tumours retain an ‘epigenetic memory’ derived from their originating cell type but also develop a tumour-specific pattern regardless of their origin. Single-cell transcriptomic analyses identified two trajectories of Club cell evolution which are similar to the ones used during lung regeneration, providing a link between lung regeneration and cancer initiation. On both routes, tumours lose their Club cell identity and gain an AT2- like phenotype. Together, this study highlights the role of Club cells in LUAD initiation and unveils key mechanisms conferring LUAD heterogeneity.

## Introduction

Neoplastic growth of non-small cell lung cancer (NSCLC) is initiated by genetic and epigenetic alterations, occurring by transforming the cell-of-origin into a (pre)neoplastic cell state. During evolution of the cancer genome, additional alterations are acquired, which often accelerate tumorigenesis. These alterations are translated into unique gene expression profiles that determine the malignant phenotype, including aggressiveness and response to therapy. The altered molecular landscapes seen in cancer, the mixture of signatures of oncogenic processes and the cell-of-origin provide an opportunity for thorough molecular characterization of malignancies and novel biomarker development (Yau et al., 2018). Recent studies on central nervous system tumours (Capper et al., 2018) and leukaemia (Lipka et al., 2017; Oakes et al., 2016; Sill et al., 2020) already harnessed this new concept for subclassification of brain tumours or outcome prediction of leukaemia therapy.

The lung epithelium is composed of diverse cell types according to their location. The upper airways include Club, Ciliated, Basal, Goblet, Neuroendocrine, and Tuft cells as well as the recently described Ionocytes (Montoro et al., 2018; Plasschaert et al., 2018). The distal airways include Alveolar type 1 (AT1) and Alveolar type 2 (AT2) cells. Many of these epithelial cells exhibit de-differentiation potential under normal homeostasis or upon lung injury. In the upper airway, Basal cells generate differentiated cells during postnatal growth (Rock et al., 2009). Club cells maintain the airway by self-proliferation and differentiation into Ciliated cells (Rawlins et al., 2009) or Basal cells, but they can also mobilize to regenerate the alveoli after damage (Kathiriya et al., 2020; Tata et al., 2013). In the distal airway, both AT1 and AT2 cells repair the injured alveoli by self-renewal and differentiation into each other (Barkauskas et al., 2013; Jain et al., 2015; Zacharias et al., 2018). Besides these well-defined epithelial cells, a rare population of double-positive CCSP/SPC Bronchioalveolar Stem Cells (BASCs), located in the terminal and respiratory bronchioles, can maintain both airway and alveoli (Liu et al., 2019; Salwig et al., 2019). Recently, single cell RNA sequencing (scRNA-seq) has contributed to the identification of a rare population of H2-K1^high^ Club-like progenitors (Kathiriya et al., 2020), the transitional stem cell state of Krt8ADI (Strunz et al., 2020), AT2-cell derived damage-associated transient progenitors (DATPs) (Choi et al., 2020), and pre-alveolar type 1 transitional cells (PATS) (Kobayashi et al., 2020), all of which contribute to lung regeneration. It remains controversial whether transdifferentiation of cells can take place during the initial transformation of normal cells, before giving rise to tumours. In this context, further characterization of this initial transition would be critical for the correct identification of the cell-of-origin of LUAD.

The cellular origin of NSCLC subgroups is largely unknown, and it is speculated that the interplay between cell types may give rise to various NSCLC subgroups. Work on genetically engineered mouse models (GEMMs) demonstrates the importance of both the cell-of-origin as well as the genetic mutation spectrum in shaping lung cancer phenotypes (Ferone et al., 2020). For instance, the cell-of-origin of *Kras*-driven LUAD has been extensively investigated using GEMMs (Desai et al., 2014; Mainardi et al., 2014; Sutherland et al., 2014; Xu et al., 2012). These studies suggest AT2 cells to be the predominant cells-of-origin of these tumours. However, rather than employing a stochastic system in which all cells have an equal chance of being transformed, these tumours were induced by forcefully expressing *Kras* in one specific cell type. Therefore, it is still highly debatable whether trans-differentiation of Club cells into AT2 cells can take place under specific environmental conditions or whether other genetic factors might alter the tumour initiating cells (Rowbotham and Kim, 2014).

More recently, we induced LUAD in adult mice by exposing them to the toxic chemicals found in tobacco smoke in combination with engineered reporters (Spella et al., 2019). We demonstrated that tobacco-induced tumours originate from airway epithelial cells which attain alveolar characteristics and not exclusively from cells originating in the alveoli. Altogether, these studies suggest that Club cells give rise to chemical-induced LUAD by attaining an AT2 phenotype, however, the developmental routes and pathways that Club cells use to become a tumour have not been investigated.

As opposed to LUAD with *Kras* mutations, the cellular origin of ALK-translocated LUAD has not yet been investigated. In the present study, we employed a stochastic LUAD model by inducing the endogenous oncogenic *Eml4-Alk* rearrangement (Maddalo et al., 2014), and took advantage of an adenovirus system that can infect any cell type within the lung epithelium. We combined the *Eml4-Alk* adenovirus with lineage-tracing mouse models, DNA methylome, and single-cell RNA transcriptome analysis and identified a subpopulation of Club cells as the cells-of-origin of *Eml4-Alk* tumours. We demonstrate that Club cells upon *Eml4-Alk* fusion trans-differentiate early during tumour development, gain the expression of alveolar markers, and follow two routes upon transformation yielding heterogeneous tumour subgroups.

## Results

### Tumour initiation and development induced by *Eml4-Alk* fusion in distinct lung cell types

To faithfully recapitulate ALK-translocated human LUAD in mice, we used a published CRISPR-Cas9 construct to induce the endogenous *Eml4-Alk* (EA) oncogenic rearrangement (Maddalo et al., 2014) (Figure 1A). In this model, intratracheal instillation of an adenovirus (hereafter Ad-EA) encoding a single guide RNA (sgRNA) targeting *Eml4*, a sgRNA targeting *Alk* and the Cas9 protein sequence, gives rise to LUAD eight weeks after instillation. To closely trace tumour initiation, we examined the lungs of multiple mice at defined time points following adenovirus delivery. At early stages of tumorigenesis, hyperplasia was found in the bronchioles and in the alveolar space. Interestingly, the bronchiolar lesions can completely engulf bronchioles, promoting partial loss of the club cell marker CCSP, while acquiring alveolar markers such as SPC (Figure 1B and 1C). Moreover, among more advanced tumour stages (adenoma and adenocarcinoma), almost every tumour was SPC positive (Supplementary Figure 1A), while rare cells inside the nodules showed traces of CCSP expression (Figure 1C).

**Figure 1.**
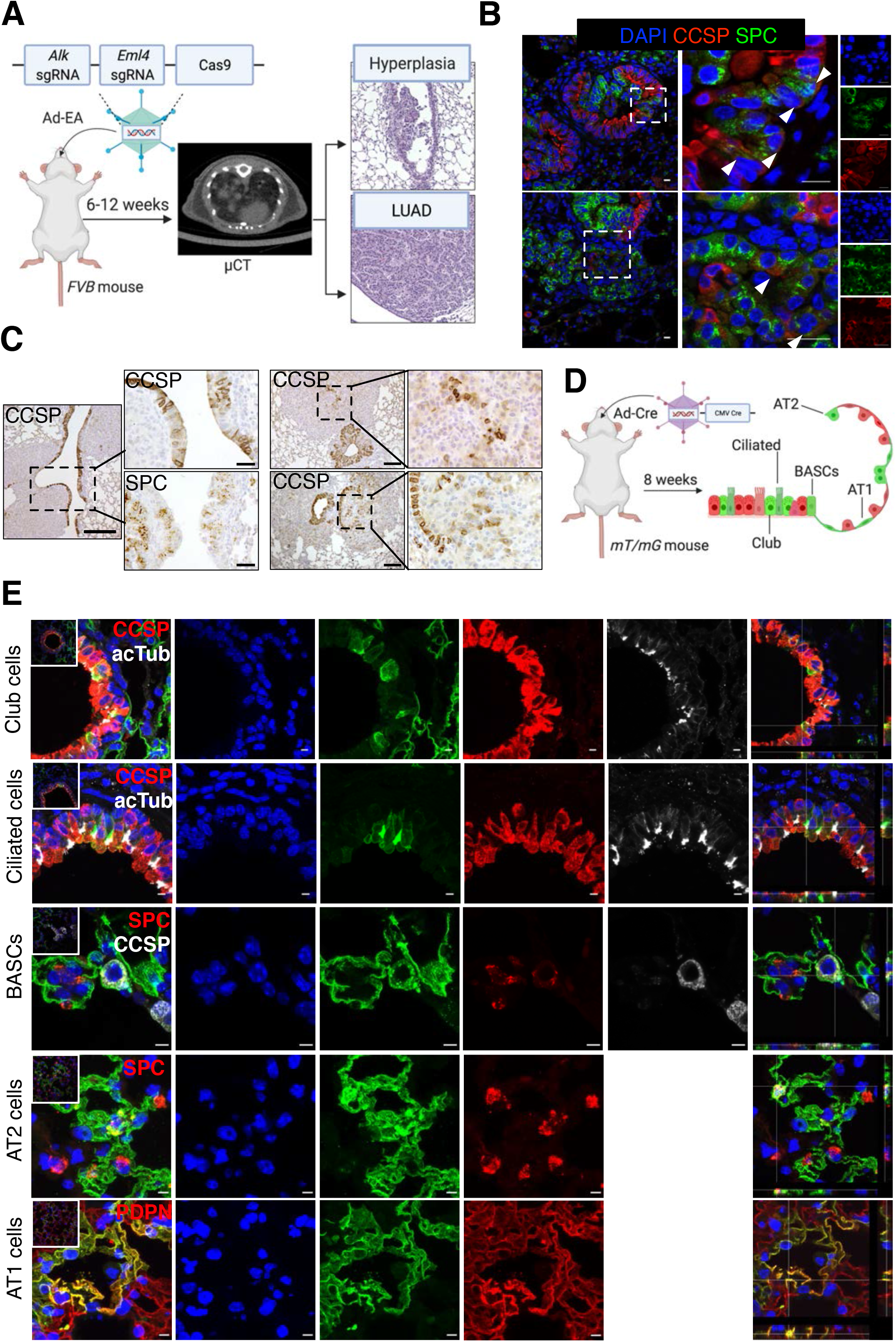
Mouse model of lung adenocarcinoma and identification of adenovirus-infected cell types. A) Schematic of the *Eml4-Alk* mouse LUAD model used. Mice were intratracheally instilled with Ad-EA and μCT was used to monitor tumour development. Representative H&E staining of an early lesion (hyperplasia) and a tumour are shown on the right. B) Immunofluorescent staining of SPC (green) and CCSP (red) antibodies showing that Club cells in the bronchi start expressing SPC upon *Eml4-Alk* rearrangement. DAPI staining in blue. Original (left), magnified overlay (middle), and their single-channel images (right) from the dashed areas are shown. Arrowheads show cells that are double-positive CCSP^+^SPC^+^. Scale bar: 20μm. C) Immunohistochemistry of CCSP and SPC antibodies from *Eml4-Alk* tumours showing CCSP^+^ bronchi as well as cells inside the tumours. Scale bars: 200μm (left panel) and 100μm (right panel). D) Experimental schematic of *mT/mG* mice transduced with Ad-Cre indicating the cell types that can get infected. E) Immunofluorescent staining of the indicated antibodies on lung sections from *mT/mG* mice transduced with Ad-Cre, showing that Club, Ciliated, BASCs, AT2 and AT1 cells are infected. Scale bars 10μm.

Compared to those mice that only received Ad5-CMV-Cre (hereafter Ad-Cre) as control, we also found an increase of double-positive CCSP/SPC cells in the bronchiolar lesions eight weeks after instillation of Ad-EA (Supplementary Figure 1B). We speculate that this increase of dual positive cells in the early lesions is a consequence of a lineage switch from Club to AT2-like cells triggered by the oncogenic transformation and not necessarily an increase in BASCs, since these cells are localized in the bronchioles and not the BADJ *(Error! Reference source not found.* Although advanced tumours uniformly expressed AT2 markers, results from early lesions suggested that other cell types, such as Club cells, might also be involved in tumorigenesis. The advantage of using an adenovirus to induce the *Eml4-Alk* rearrangement is that any cell type within the lung epithelium has the chance of being infected. To define the cell tropism of the adenovirus, we used Ad-Cre to instil a mouse strain that switches membranous tdTomato to membranous EGFP fluorescence *(mT/mG)* (Muzumdar et al., 2007) upon Cre mediated recombination. We identified that virtually all lung epithelial cell types got infected with Ad-Cre (Figure 1D, 1E and Supplementary Figure 1D), being AT1 cells the largest cell population infected, followed by AT2, Club, and Ciliated cells. We rarely found infected BASCs possibly due to their low frequency compared to the other cell types (Supplementary Figure 1E).

From these results, we hypothesize that the *Eml4-Alk* oncogenic rearrangement can initiate tumours in Club and AT2 cells, although late-stage tumours homogeneously express the AT2 cell marker, SPC.

### Club and AT2 cells give rise to Eml4-Alk lung adenocarcinomas

To evaluate the cell-of-origin of *Eml4-Alk* induced LUAD, we crossed *mT/mG* mice with *Scgbla1-CreERT* (Rawlins et al., 2009) for labelling Club cells (hereafter *Scgb1a1), Sftpc- CreERT2-rtTA* (Chapman et al., 2011) for AT2 cells (hereafter *Sftpc), Hopx-CreER* (Takeda et al., 2011) for AT1 cells (hereafter *Hopx*) and *Foxj1-CreER* (Rawlins et al., 2007) for Ciliated cells (hereafter *Foxj1*). Next, we assessed the specificity of each lineage-tracing mouse line, by performing immunohistochemical and immunofluorescent staining of the corresponding cell markers together with GFP after tamoxifen-induced (TAM) recombination (Figure 2A and Supplementary Figure 2A). In line with previous reports (Kathiriya et al., 2020), the *Scgb1a1* line labelled not only Club cells, but also AT2 and Ciliated cells (Figure 2B). Similarly, the *Hopx* line labelled Club and Ciliated cells in addition to AT1 cells (Figure 2A and 2B). However, unspecific labelling was not found in *Sftpc* and *Foxj1* lines.

**Figure 2.**
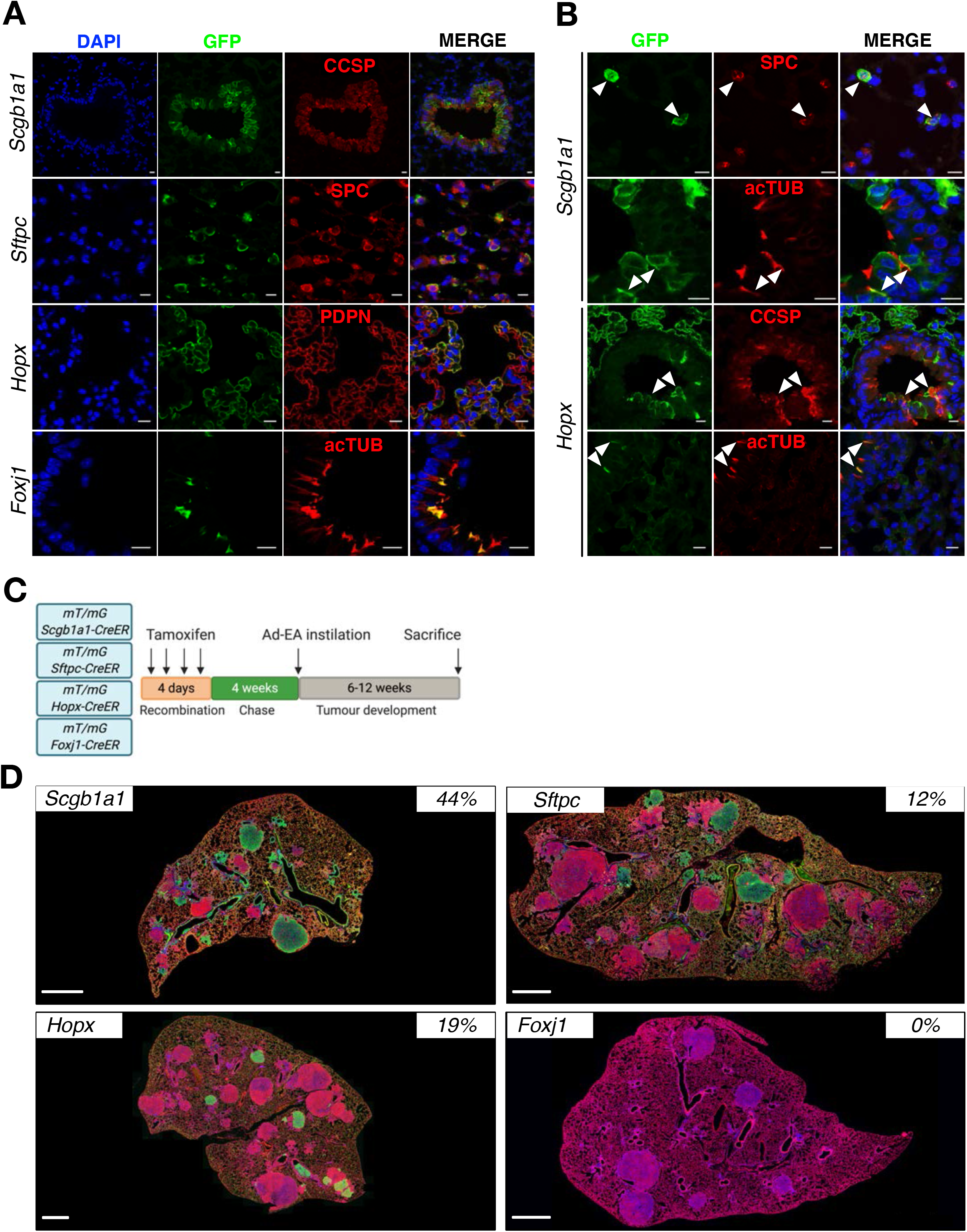
Lineage-tracing models. A) Immunofluorescent staining of GFP and the indicated antibodies in the different lineage-tracing mouse models (*Scgb1a1, Sftpc, Hopx and Foxj1*) showing the specific labelling (CCSP, SPC, PDPN and acTUB). Images of single-channel and overlay are displayed sequentially. Scale bars: 10 μm. B) Immunofluorescent staining of GFP and the indicated antibodies (CCSP, SPC and acTUB) in the lung sections from lineage-tracing mouse models showing the labelling specificity. Arrows indicate the unspecific labelling of the cells. Scale bars: 10 μm. C) Schematic of the labelling and tumour induction of linage-tracing mice. D) Immunofluorescent staining of GFP and RFP antibodies in the respective mouse models. The percentages on the upper right corners represent the number of green labelled tumours out of the total number of tumours analysed in each line. Scale bars: 500μm.

Subsequently, we induced LUAD in the lineage-tracing mice by intratracheal instillation of Ad-EA four weeks after TAM injection. When tumours became discernible by μCT, we analysed the percentage of GFP^+^ tumours in each line (Figure 2C and 2D). We found that *Scgb1a1* mice had a higher rate of GFP^+^ lesions (44%±5) than *Sftpc* (12%±9) (Figure 2D and Supplementary Figure 2B). Surprisingly, almost one-fifth of the tumours in *Hopx* mice were also GFP^+^ (19%±14), although AT1 cells were rarely reported to contribute to LUAD (Jain et al., 2015). Due to labelling promiscuity of the *Hopx* line, we further tested the possibility that these GFP^+^ tumours were arising from Club rather than AT1 cells. Therefore, we infected *mT/mG; Krt5-CreER* mice (hereafter *Krt5*) (Indra et al., 1999) with Ad-EA, since only AT1 cells in the distal lung, besides Basal cells in the trachea, are labelled in this model (Figure 3A and B and Supplementary Figure 3A). However, we failed to observe any GFP^+^ tumour in *Krt5* mice *(0 out of 130 analysed tumours),* suggesting that neither Basal cells from upper airways nor AT1 cells contributed to *Eml4-Alk* LUAD initiation (Figure 3C and Supplementary Figure 3B). Therefore, the GFP^+^ tumours arising from the *Hopx* line most probably originated from unspecifically labelled Club cells (Figure 3D and E). In addition, all tumours developed in *Foxj1* mice were negative for GFP (*0 out of 113 analysed tumours)* (Figure 2D and Supplementary Figure 2B). Notably, the GFP^+^ tumours from *Hopx, Scgb1a1,* and *Sftpc* lines exhibited the same expression pattern: positive for SPC but negative for CCSP (Figure 3E and Supplementary Figure 3C).

**Figure 3.**
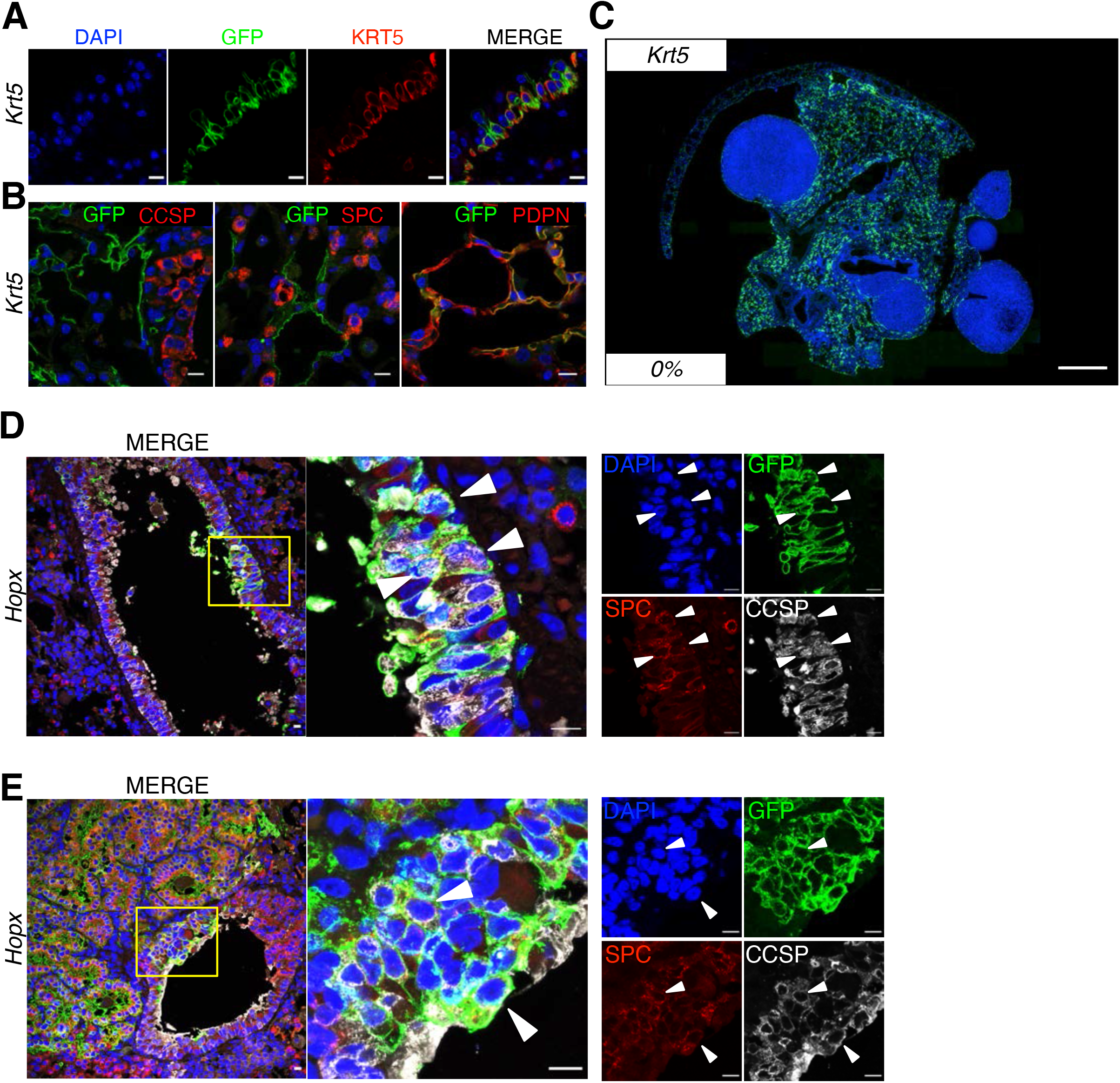
Characterization of Krt5 and Hopx mice. A) Immunofluorescent staining of GFP and KRT5 antibodies in the trachea of *Krt5* mice showing labelled Basal cells. Overlay and single-channel images are sequentially presented. Scale bars: 10μm. B) Immunofluorescent staining of GFP and the indicated antibodies in the distal lung of *Krt5* mice. AT1 cells, but not Club or AT2, are labelled. Blue (DAPI), green (GFP), red (as indicated). Scale bars: 10μm. C) Overview of GFP immunofluorescent staining in *Krt5* mice. The percentage on the lower left corner represents the percentage of GFP^+^ tumours. Scale bars: 1mm. D, E) GFP, SPC, and CCSP immunofluorescent staining on *Hopx* mice trasnduced with Ad- EA. Both early (D) and late (E) stages of tumorigenesis are shown; the original, magnified and single-channel images are sequentially shown from left to right; arrowheads indicate the labelled Club cells under lineage switch into AT2 cells. Scale bars: 10μm.

Taken together, these findings establish both Club and AT2 cells as the cell-of-origin of *Eml4-Alk* induced LUAD, although Club cells seem to be the main contributors. Additionally, the phenotypic similarity of the tumours suggests that Club cells might undergo a lineage switch upon oncogenic transformation.

### Lung cancer and cell type specific DNA methylation patterns

Epigenetic reprogramming during differentiation is tightly regulated in a lineage-specific manner (Oakes et al., 2016), resulting in cell type-specific DNA methylation (Dor and Cedar, 2018). Patterns of different developmental routes are preserved as ‘epigenetic memories’ within each cell type (Kim and Costello, 2017). Therefore, we sought to uncover signatures of each specific cell type originated tumour and to trace them back to their cells-of-origin. We captured the DNA methylation landscape of sorted GFP^+^ cell populations from lineage-labelled healthy mice and tumours from the *Sftpc, Hopx,* and *Scgb1a1* lines (Supplementary Figure 4A) by tagmentation-based whole-genome bisulfite sequencing (TWGBS) (Wang et al., 2013).

Using TWGBS data, we determined regions which could distinguish different cell types based on DNA methylation differences. We found that poised enhancers, as opposed to active enhancers and promoters, were specifically powerful discriminating normal lung cell types. However, in principal component analysis (PCA), tumours showed homogenous methylation patterns and did not cluster according to their cellular origin, as did the normal cell types (Figure 4A and Supplementary Figure 4B). To infer cell type composition, we applied a reference-free deconvolution method, MeDeCom, to the DNA methylation data of the most variable CpG sites overlapping with bivalent enhancers. MeDeCom identifies so-called latent methylation components (LMCs) designed to capture the proportion of cell type-specific methylation signatures in each sample (Lutsik et al., 2017; Scherer et al., 2020).

**Figure 4.**
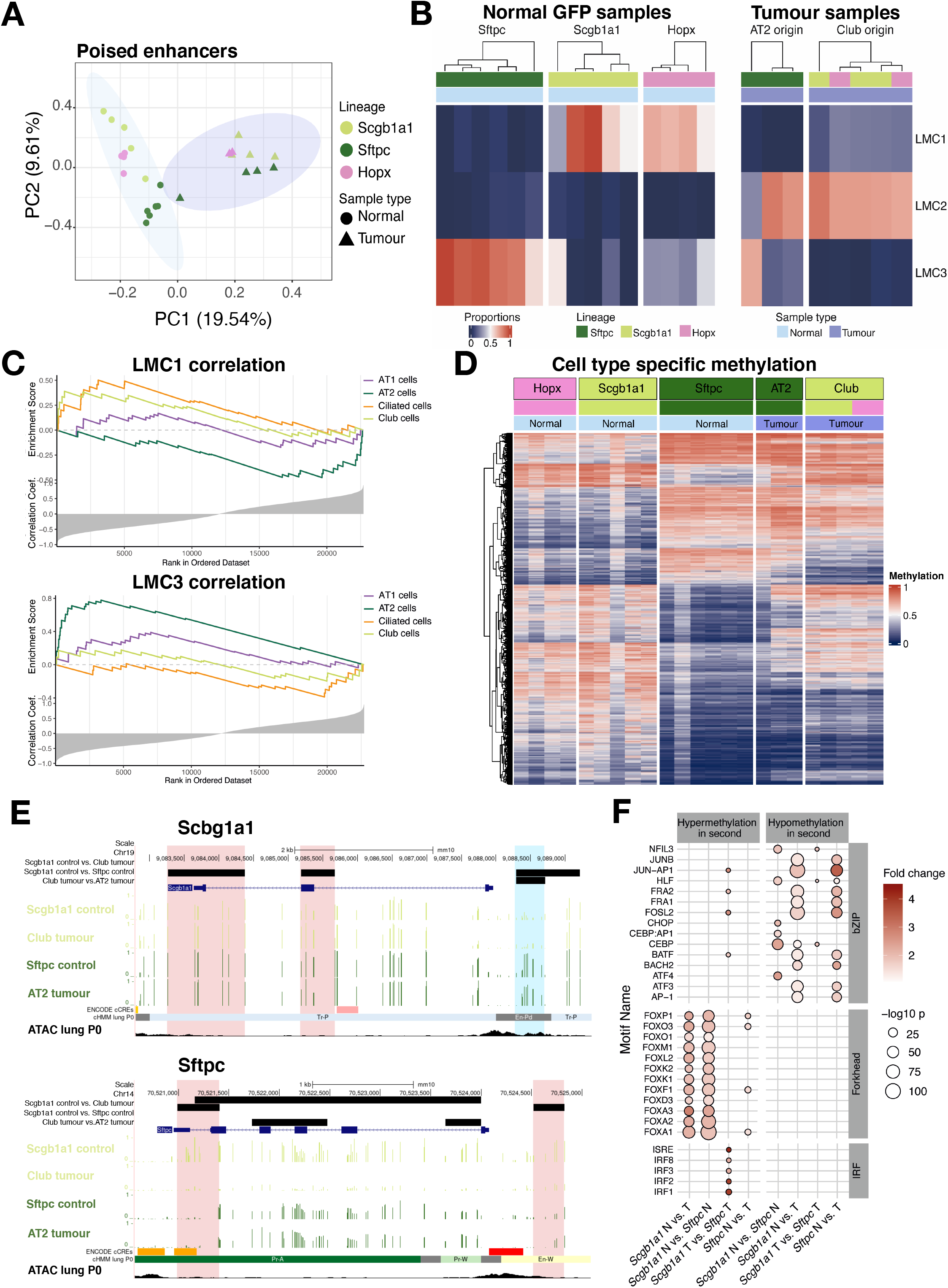
DNA methylation patterns. A) Principal component analysis of DNA methylation at TSS-distal and TSS proximal poised enhancer regions (En-Pd and En-Pp, based on ENCODE postnatal 0 days mouse lung, ENCSR538YJF). The shape of the dots reflects the sample type, as normal lung (Normal) or GFP^+^ tumours (Tumour). The colour indicates mouse lineages used. Coloured shading is drawn around the Normal (light blue) and Tumour (purple) samples. B) DNA methylation deconvolution by MeDeCom analysis shows three identified latent methylation components (LMC 1-3). Colours indicate the proportion of LMCs for each sample. Samples are clustered within each mouse lineage *(Sftpc, Scgb1a1* or *Hopx)* and according to sample type (Normal/Tumour). C) Association of LMCs with cell type-specific gene signatures based on published singlecell RNA sequencing data (Strunz et al., 2020). Genes are ranked by their correlation of promoter methylation with the respective LMC. Enrichment scores are running sums calculated using the Gene Set Enrichment Analysis algorithm. D) Heatmap of regions that were differentially methylated between *Scgb1a1* and *Sftpc* normal samples. The methylation value is shown as beta values ranging from 0 to 1 visualized by the heatmap colours blue to red. For tumours, the label of each column shows the proposed originating cell type, while the colours below depict their originating lineage. E) Average methylation across the *Scgb1a1* and *Sftpc* genes for normal samples from *Scgb1a1* and *Sftpc* lines and Club and AT2 originated tumours. Pink bars highlight regions with DNA methylation differences between normal lineages, while blue bars highlight regions with differences in tumours from distinct origins. ENCODE cCRE, showing candidate Cis-Regulatory elements, cHMM lung P0 showing ChromHMM regions in mouse lung, postnatal 0 days, and ATAC lung P0 showing ATAC-seq peaks in postnatal 0 day old lung are UCSC tracks under the same name. F) TF motif enrichment analysis of the differentially methylated regions (DMRs). Each column represents one comparison (e.g. *Scgb1a1* normal vs. tumour) in one direction (e.g. hypermethylation in second, as in hypermethylated in tumours). The colour of the dots shows the enrichment of the DMRs for the motifs compared to random genomic regions. The size of the dots reflects on the –logιo(p value) of the enrichment analysis. Empty lines show the lack of significant enrichment.

After optimization and quality control of the methylome deconvolution model (for details, see methods), we identified three main LMCs: LMC1 and LMC3 represented normal samples from *Scgb1a1* and *Sftpc* lineages, respectively, whereas LMC2 represented a common lung cancer-specific signature (Figure 4B). We could not identify a specific component for AT1 (appearing in *Hopx* lineage) or Ciliated cells (appearing in *Scgb1a1* lineage). To define the cell types represented by each LMC, we performed an enrichment analysis of the cell typespecific markers, previously identified by scRNA-seq (Han et al., 2018; Strunz et al., 2020; Travaglini et al., 2019). Since promoter methylation is considered to correlate negatively with gene expression, the enriched gene signatures captured the cell types associated with the given component. We therefore, correlated the DNA methylation levels of each promoter with the proportion of each LMC and determined whether cell type-specific markers were overrepresented among the inversely correlating genes (Figure 4C). The results revealed that LMC1 represented not only Club but also Ciliated cells. Similarly, LMC3 captured both AT2 and AT1 cells (Figure 4C). Importantly, tumour samples, despite showing a homogeneous pattern in LMC2, still resembled the specific signature of their cell-of-origin, classified by a higher contribution (>10%) of the respective components (Figure 4B). A high proportion (>10%) of LMC1 in both tumours from the *Hopx* line supported our previous result that these tumours are derived from Club cells.

Differential methylation analysis revealed cell type-specific differences between AT2 and Club cells. Notably, tumours, irrespective of their cell-of-origin, showed AT2-like methylation patterns (Figure 4D). These results strengthened the assumption that Club cell originating tumours switch their identity during tumorigenesis. This was also supported by the promoter methylation patterns of *Sftpc* and *Scgb1a1* genes (Figure 4E), which showed that irrespective of their origin, the tumours showed patterns similar to those found in normal AT2 cells. To gain mechanistic insight into the lineage switch of Club cells during tumorigenesis, we analysed the enrichment of transcription factor (TF) binding motifs in differentially methylated regions (DMRs) (Figure 4F and Supplementary Table 1). We observed strong enrichment of Forkhead family TFs among regions that were hypermethylated in Club cell originating tumours, as well as in normal AT2 cells compared to normal Club cells. Forkhead TFs regulate lung secretory epithelial fate (Li et al., 2012) and play a role in maintaining epithelial cell identity (Paranjapye et al., 2020). Therefore, increased methylation of their binding site might contribute to derailing the normal lung regeneration processes. Tumour-specific hypomethylation events were enriched in TF binding motifs of the JUN/FOS proteins, building blocks of the activator protein-1 (AP-1) transcription factor known for their role in lung tumorigenesis as well as their involvement within other signal transduction pathways (Karamouzis et al., 2007).

Taken together, the DNA methylation data supports the Club cell origin of labelled tumours from *Scgb1a1* and *Hopx* lines. Furthermore, the DNA methylation patterns and differential methylation of TF binding motifs suggest an epigenetically-driven lineage switch of Club cells into an AT2-like phenotype before or during tumorigenesis.

### Single-cell RNA sequencing identifies multiple cell states after transformation of Club cells

Lineage-tracing mouse models and DNA methylation data indicated that tumours originating from Club cells acquire an AT2-like phenotype. To elucidate the molecular processes of tumorigenesis, we designed a time-course experiment based on scRNA-seq of the *Scgb1a1* line (Figure 5A). We collected two control samples; one with only TAM injection and another one 18 weeks after TAM and Ad-Cas9 infection (Ad-Cas9); two early samples 2 weeks after Ad-EA and one intermediate sample 4 weeks after Ad-EA. We isolated live Club cells (EpCAM^+^/ CD45^-^/ CD31^-^/ tdTomato^-^/ GFP^+^/ CD24^-^/ ß4^+^/ CD200^+^) from these samples (Figure 5A and Supplementary Figure 5A). To discern possible implications of the labelled AT2 cells in the *Scgb1a1* model, we included all GFP^+^ cells from a second intermediate sample - 4 weeks after Ad-EA - as well as all GFP^+^ tumour cells from late-stage tumour nodules (Figure 5A and Supplementary Figure 5A) and performed scRNA-seq.

**Figure 5.**
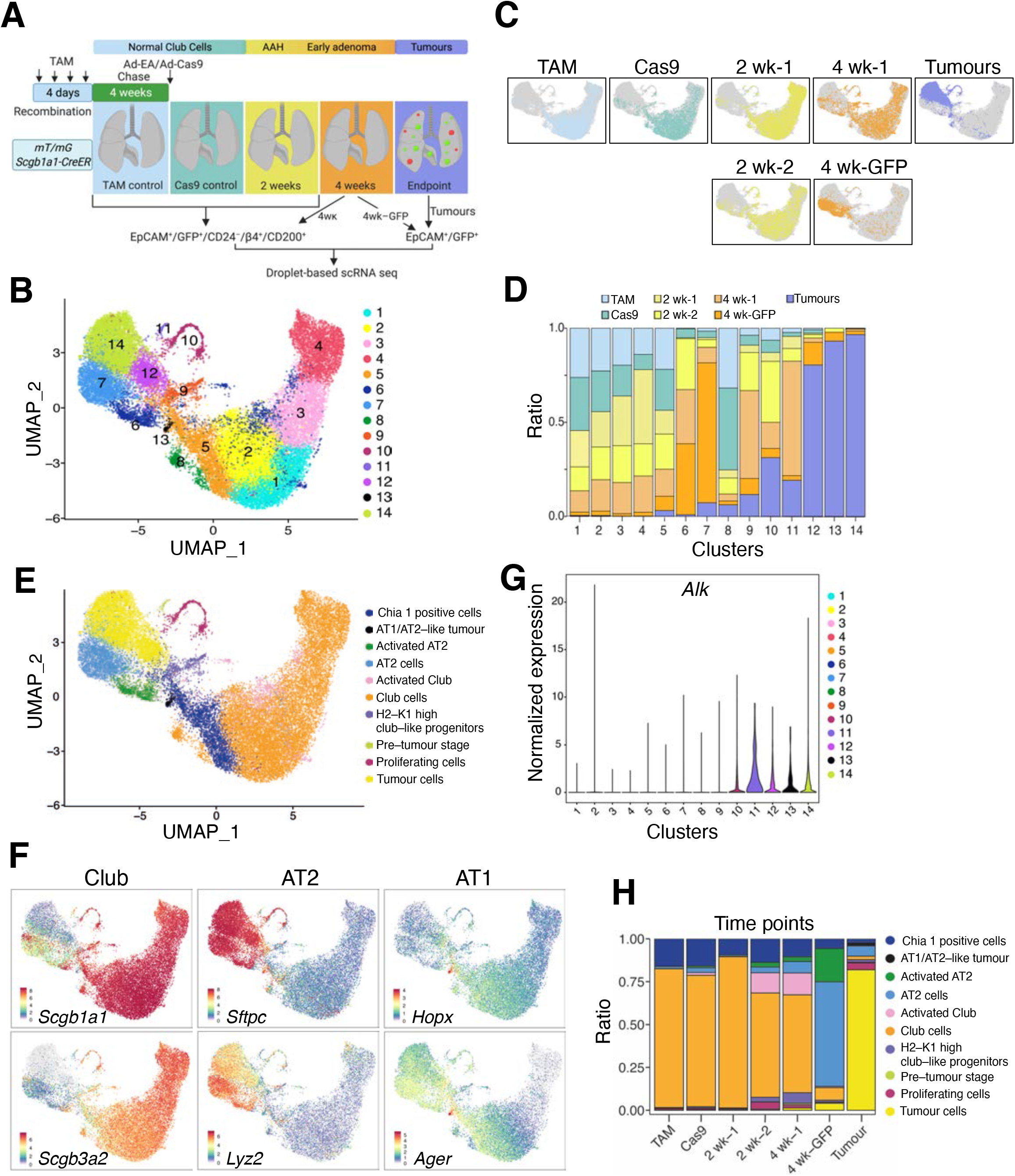
Single-cell RNA sequencing. A) Experimental strategy of the scRNA sequencing analysis. B) UMAP of the high-quality cells. Cells are coloured by the clusters identified using a shared nearest neighbour (SNN) modularity optimization-based clustering. C) UMAP embedding showing the distribution of the cells collected from the different time points. Coloured dots represent the cells collected at the indicated time point; grey dots represent cells from the other timepoints. D) Percentage of cells in each cluster, coloured by sample origin. The relative contributions were normalized to the number of cells in each sample. E) UMAP coloured by the cell type annotation of the study set. The annotation combines previously published cell type specific markers (Angelidis et al., 2019; Han et al., 2018) and manual curation and is based on the most significant marker genes of each cluster or cell group (Supplementary Table 3). F) UMAP with cells coloured by expression of Club *(Scgb1a1, Scgb3a2),* AT2 *(Sftpc, Lyz2)* and AT1 (*Hopx, Ager*) cell type-specific markers. Colours represent the normalized expression levels for each marker in each cell. G) Violin plot depicting the normalized expression of *Alk* by cluster indicating elevated expression in clusters 11, 12, 13, and 14. H) Cell type composition of each collected sample.

Unsupervised cell-clustering analysis of 29,277 high-quality cells uncovered 14 unique clusters (Figure 5B and Supplementary Table 2), showing a distinct distribution of the samples throughout tumour progression (Figure 5C and D). As indicated in Figure 5D, clusters 1, 2, 3, 4, 5 and 8 were mainly (22-75%) composed of TAM and/or Cas9 control cells, while clusters 12, 13 and 14 were predominantly (>80%) composed of tumour cells. Clusters 6, 9, 10 and 11 were mainly composed (>55%) of cells derived from the 2 and 4 weeks transduced animals (Figure 5C and D). Notably, the majority (74%) of cluster 7 came from the 4 weeks GFP sample, where labelled normal AT2 cells had not been excluded.

Next, clusters were annotated using cell type-specific markers (Angelidis et al., 2019; Han et al., 2018) and previously published cell type signatures (Supplementary Table 3) (Kathiriya et al., 2020; Marjanovic et al., 2020; Strunz et al., 2020). As expected, multiple clusters were annotated as Club cells and in line with our previous results, tumour clusters presented high levels of AT2 markers, except for cluster 13, which unexpectedly showed increased levels of AT1 markers (Figure 5E, 5F, and Supplementary Table 3). Furthermore, due to the high expression of *AW112010, H2-K1,* and *Cd74,* we identified cluster 9 to be similar to the recently described H2-K1^high^ Club-like progenitor cells which were shown to contribute to lung regeneration after injury (Choi et al., 2020; Kathiriya et al., 2020; Kobayashi et al., 2020) (Supplementary Figure 5B and Supplementary Table 3).

To verify whether tumour cells contained the *Eml4-Alk* rearrangement, we checked for the aberrant expression of the *Alk* region affected by the translocation. We found *Alk* expression not only in the tumour clusters 12, 13 and 14 but also in cluster 11 (Figure 5G). Moreover, we observed a progressive loss of Club cell identity and gain of AT2-like features throughout tumour development (Supplementary Figure 5C). Cluster 11 still showed a Club signature and low expression of tumour markers (Supplementary Figure 5D and 5E), suggesting its pre-tumour stage.

Collectively, our scRNA-seq analyses identified distinct cell populations that were enriched in different stages of Club cell tumorigenesis. A dynamic change of cell type composition was observed with the rise of progenitor-like cell states in early time points and distinct tumour clusters being discernible as early as 4 weeks after oncogenic induction (Figure 5H).

### Club cells follow two routes towards LUAD

We next sought to model the differentiation trajectory of Club cells towards LUAD. RNA velocity (La Manno et al., 2018) estimates the ratio of spliced and unspliced mRNA predicting the future state of cells. Partition-based graph abstraction (PAGA) analysis (Wolf et al., 2019) generates a map, where nodes are connected by weighted edges representing the connectivity between clusters. Using RNA velocity (Supplementary Figure 6A) to direct the PAGA edges (Supplementary Figure 6B), we obtained an unbiased estimation of linage trajectories (Bergen et al., 2020; Wolf et al., 2019) (Figure 6A).

**Figure 6.**
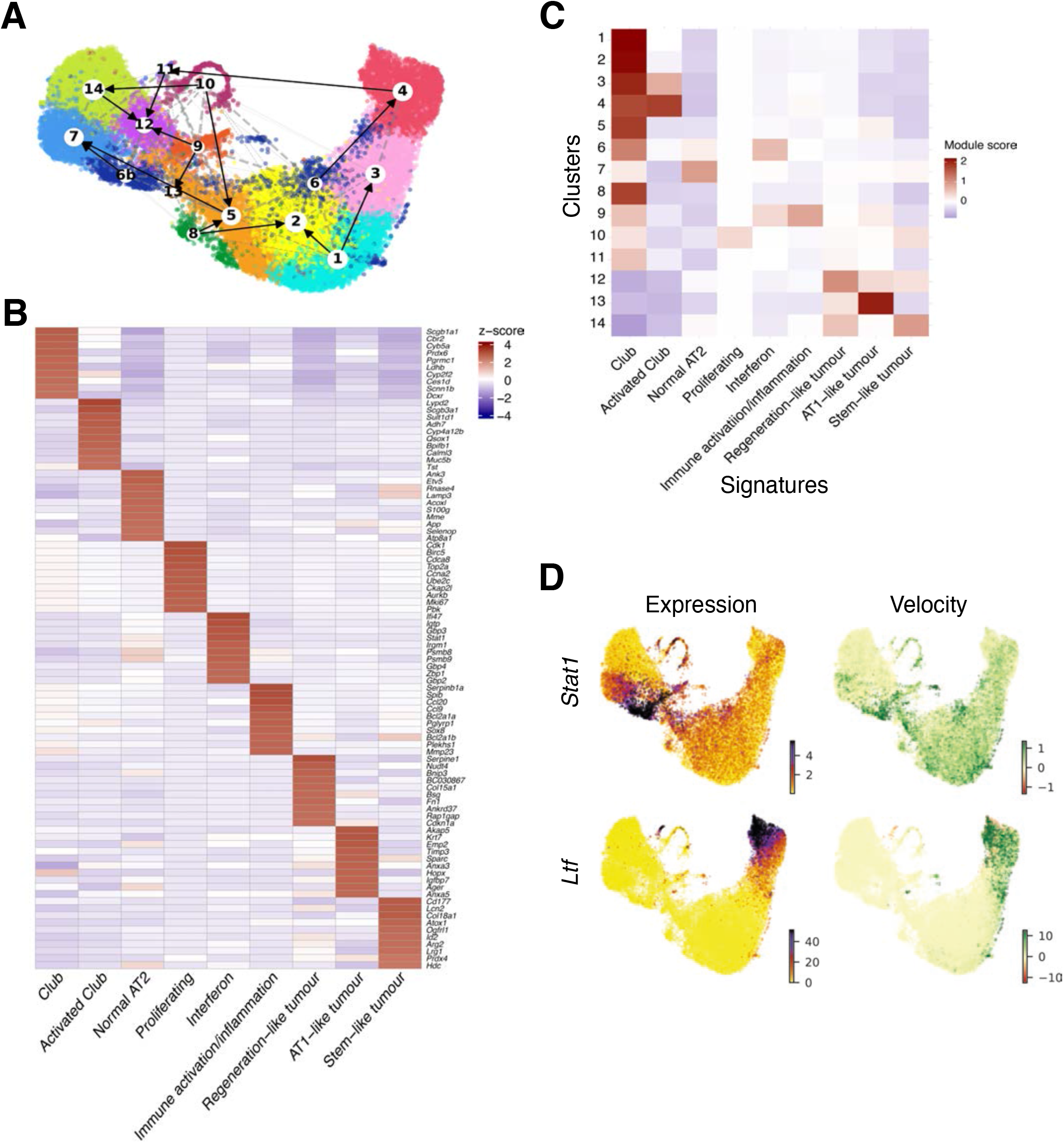
Activity programs. A) UMAP embedding of combined RNA velocity and PAGA. Connections between clusters are established based on PAGA, while the direction of the arrows is inferred from the RNA velocity analysis. B) Heatmap of the module scores of each cluster (Y axis) for the identified activity programs (cNMF modules, STAR Methods). The top 10 genes were used to calculate the module score. C) Heatmap displaying the top 10 marker genes of each activity program identified by the cNMF analysis. The heatmap colours show the z-score unit gene expression in each activity program. D) RNA velocity and gene expression of candidate genes *Stat1* (upper) and *Ltf* (lower). The right side of the plots shows the RNA velocity. The green colour represents a high velocity, therefore upregulation, while red colour denotes downregulation. The left side of the plots shows the current gene expression levels.

To better understand the transitions of Club cell states during tumour progression, we explored gene expression activity programs using consensus non-negative matrix factorization (cNMF) (Kotliar et al., 2019; Marjanovic et al., 2020). We identified nine activity programs (Figure 6B and Supplementary Table 4), three of which depicted the tumour clusters (12, 13, and 14) described above: a stem-like tumour module correlated with cluster 14, and was characterized by stem cell markers such as *Id2* and *Sox9* together with AT2 identity genes like *Sftpc, Lamp3* and *Lcn2* (Figure 6B, 6C and Supplementary Table 4); a regeneration-like tumour module that was mainly enriched in cluster 12 and similar to lung regeneration, this module showed high activity in HIF1 signalling, TGF-beta pathway, IL-17 signalling and metabolic pathways such as glycolysis (Supplementary Figure 6C and Supplementary Table 4) (Choi et al., 2020). The third one, an AT1-like tumour program was highly active in cluster 13, although also present in cluster 12 (Figure 6C). This module showed high similarity to recently described signatures implicated in lung regeneration (Choi et al., 2020; Kobayashi et al., 2020; Strunz et al., 2020) as well as to “mixed AT1/AT2” and “highly mixed” programs described in AT2 originated *Kras* mutant lung tumours (Marjanovic et al., 2020) (Supplementary Figure 6D, 6E and Supplementary Table 3). These results suggest that *Eml4-Alk* Club cell originated tumours are highly heterogeneous, similar to the *Kras* AT2 originated ones (Marjanovic et al., 2020).

The intermediate progenitor-like states (clusters 6 and 9) showed high activity of two immune-related programs: an interferon program and an immune activation/inflammation program (Figure 6C and Supplementary Table 4). Since cluster 6 consisted of two separated cell types, we split the cluster into two for further analysis: 6 for the Activated Club cells and 6b for the Activated AT2s. RNA velocity (Supplementary Figure 6A), and PAGA analysis (Supplementary Figure 6B) predicted clusters 6 and 9 to be connected to the tumour clusters. Therefore, we hypothesize that these progenitor cells could represent a crucial state in determining the fate of Club cell originated tumours.

Based on these results, we postulated that progenitor-like cells (clusters 6 and 9) with an active interferon signalling can follow two major routes towards tumorigenesis. The first one through proliferating cells (cluster 10), before giving rise to tumour cells (cluster 14). The second route underwent the pre-tumour state (cluster 11) and further differentiated into tumour cells in clusters 12 and 13, with the latter also connected with cluster 9. Notably, the second route was also partly shared by a subpopulation of Club cells in cluster 4 that showed a Goblet-like gene signature (upregulated *Ltf, Bpifa1, Bpifb1, Reg3g* expression), and differentiated into cluster 11 (Supplementary Table 2). Our results show that although different tumour clusters arise on diverse paths, these paths are interconnected. As representatives of these routes, we selected two candidate genes. *Stat1*, a marker of the interferon program, was upregulated mainly in the progenitor clusters, as inferred from expression and RNA velocity (Figure 6D). *Ltf,* a key player of the second route, was highly expressed in clusters 4 and 11, although RNA velocity showed spliced transcripts in cluster 11 predicting its downregulation (Figure 6D).

To further confirm these tumorigenic routes, we checked the expression of *Stat1* and *Ltf* in lung sections of different stages of tumour development (Figure 7A and B). In agreement with our computational analysis, we identified both nuclear *Stat1* and *Ltf* expression in Club cells in early lesions, but rarely in normal Club cells or tumour cells. Interestingly not all bronchi were positive for *Stat1*, implying that either those cells were not transduced by EA or that these cells followed a different route. These results validate the activation of interferon signalling (*Stat1*) in Club-like progenitor cells, as well as the involvement of Club cells with Goblet-like *(Ltf)* signature in tumour progression (Figure 7C).

**Figure 7.**
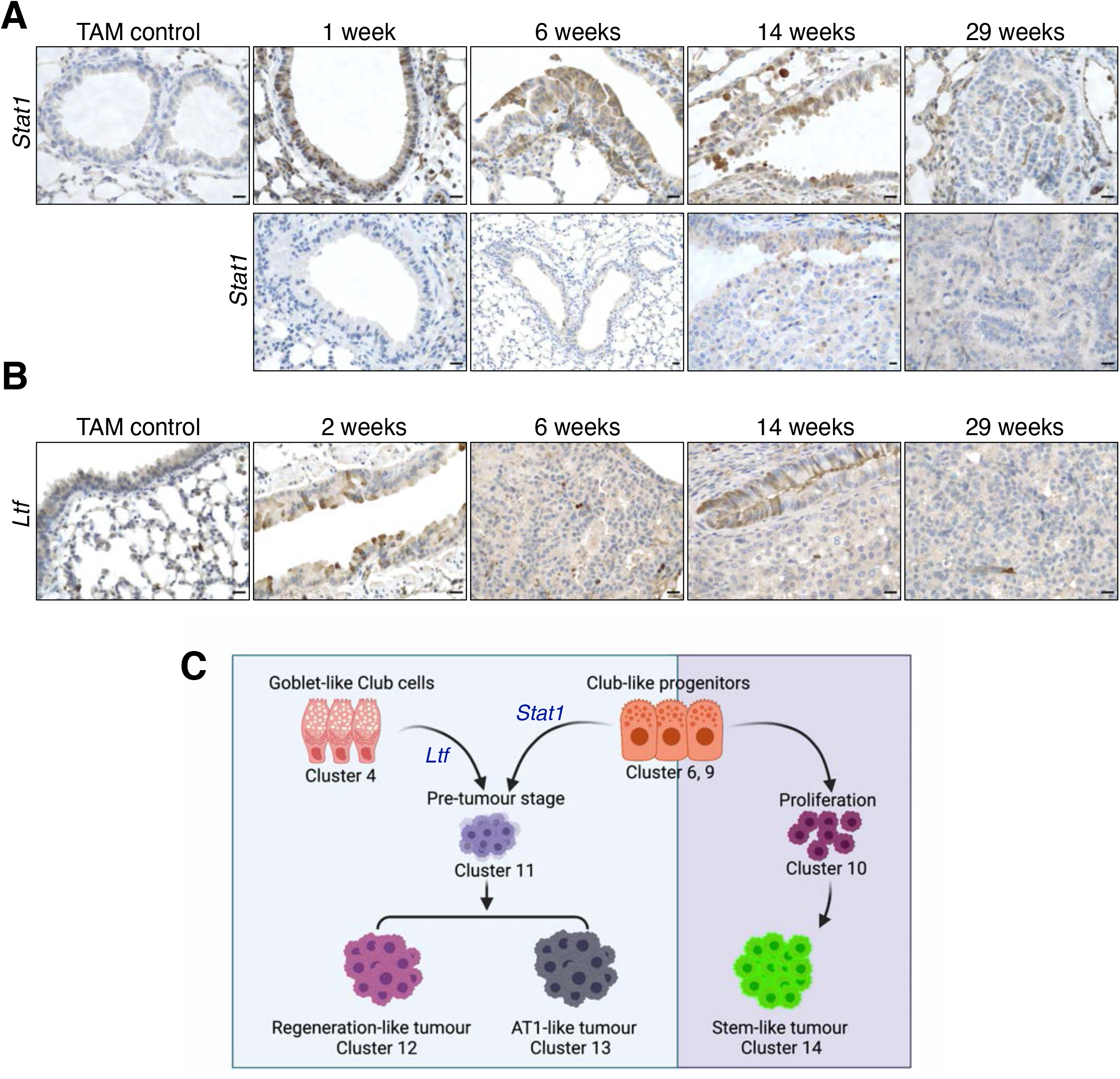
Validation of the tumorigenic routes. A) Immunohistochemistry of STAT1 in TAM control samples and in representative *Eml4-Alk* lung sections at different time points after adenoviral instillation. Upper panel shows positive staining in some bronchi at early stages while lower panel shows no staining in others. Scale bar 20 μm. B) Immunohistochemistry of LTF in TAM control samples and in representative Eml4-Alk lung sections at different time points after adenoviral instillation. Scale bar 20μm. C) Schematic representation of the two routes followed by Club cells upon *Eml4-Alk* transformation.

## Discussion

The identification of the cell-of-origin of lung adenocarcinoma is a long-standing question in cancer research. Here, we combined state-of-the-art lineage-tracing mouse models of lung cancer, DNA methylome and single-cell transcriptome analysis and showed that due to the high plasticity of lung epithelial cells, tumour characteristics are only partly dependent on their originating cell type. Previous studies focused on defining the cellular origin of LUAD, by using *Kras* mutant models that were based on tumour induction in specific cell types (Kim et al., 2005; Mainardi et al., 2014; Spella et al., 2019; Sutherland et al., 2014; Xu et al., 2012). Beyond these well-stablished *Kras-*lung cancer models, the cell-of-origin of LUAD in cell type independent oncogene-driven LUAD is largely unknown. Here, we employed lineage-tracing mouse models to label Club, AT2, AT1, Ciliated and Basal cells and induced LUAD by a viral instillation that generates an *Eml4-Alk* oncogenic rearrangement in lung epithelial cells. We identified Club and AT2 cells as the main cell types implicated in LUAD development, and showed that *Eml4-Alk* fusion gene in AT1, Ciliated or Basal cells did not give rise to tumours.

Remarkably, regardless of the originating cell type, all *Eml4-Alk* tumours were positive for the AT2 cell marker, SPC, indicating a possible lineage switch of Club cells during tumour progression. In line with this observation, Sutherland et al. previously showed that CCSP^+^ *Kras* mutant hyperplastic cells gradually lose the expression of CCSP and gain expression of SPC (Sutherland et al., 2014). DNA methylation was shown to act as a cellular memory, storing information on previous differentiation states (Kim and Costello, 2017). After deconvolution of the DNA methylomes from Club and AT2 originating tumours, cell type-dependent signatures were retained despite the large similarity between tumours. Globally, and at certain genes, tumour methylation independent of their origin, was similar to that in AT2 cells, further supporting that Club cells switch their cellular identity and rewire their epigenomic landscape during tumorigenesis.

Our findings are especially interesting in comparison with cell-of-origin studies in haematological cancers. Normal haematopoiesis is characterized by unidirectional methylation changes(Oakes et al., 2016; Touzart et al., 2021) and leukaemia cells preserve large parts of these epigenomes (Wierzbinska et al., 2020). This is in contrast to the patterns seen in LUAD where a dynamic, network-like differentiation landscape establishes highly similar tumour methylomes, arising on diverse pathways. Remarkably, the DNA methylation patterns in both cases can be used to infer the cell-of-origin.

Our scRNA-seq analysis of Club cells throughout tumour development revealed an expansion of progenitor-like cells at early stages of tumorigenesis. We observed high heterogeneity and phenotypic diversity, similar to AT2-originated *Kras* mutant tumours (Marjanovic et al., 2020). We identified three different tumour-related transcriptional programs. We speculate that since the endpoint tumour sample consisted of multiple tumours pooled from the same animal, it is possible that these tumours were in different stages or even obtained different progression patterns.

Tumorigenesis can occur as a disruption of normal regeneration processes, as recently described in pancreatic cancer (Alonso-Curbelo et al., 2021). Club cells are able to regenerate the lung by renewing their own population (Zheng et al., 2017) and transdifferentiating into AT2 and AT1 cells. Similar to lung regeneration, this high plasticity of Club cells was observed during *Eml4*-*Alk* tumorigenesis, where inflammatory and environmental stimuli rewired the transcriptome and epigenome of Club cells to evolve to an alternative fate program.

Interestingly, the tumorigenic routes followed by Club cells show certain correlation to some activity programs previously reported in AT2 originated tumours (Dost et al., 2020; LaFave et al., 2020; Marjanovic et al., 2020). This suggests that although lung tumours can have different cells-of-origin they can converge into similar stages of tumorigenesis.

By integrating scRNA-seq with whole-genome bisulfite sequencing analysis of *Eml4-Alk* tumours, we show that LUADs, unlike other tumour types, are only partly dependent on their originating cell type. In contrast, they are primarily determined by the oncogenic signals and further signalling pathways that are hijacked by the tumour cells. We identified that *Stat1* and *Ltf* shape cell differentiation throughout cancer progression and evoke signalling cascades that frequently govern normal lung development and regeneration, with direct implications in the route the neoplastic lesions will follow during tumorigenesis. We describe the paths that Club cells follow resulting in a highly heterogenous state through two major tumorigenic routes. Characterisation of these mechanisms, which are shared or unique in the different cancer subtypes, will help to decipher the routes of lung cancer initiation and identify actionable targets for personalised treatment of LUAD.

## Acknowledgments

We are grateful to Severio Bellusci and Harold Chapman for providing *Sftpc-CreER* mice and to Claudia Sholl and Brigit Hogan for *Scgb1a1-CreER* and *Foxj1-CreER* mice. We thank Simone Kraut, Marion Bähr, the DKFZ Core Facilities of Light Microscopy, Flow Cytometry, Small Animal Imaging Center and Genomics and Proteomics for the excellent technical assistance; and the Central Animal Laboratory for animal husbandry. We appreciate the help of Jan-Philipp Mallm and the DKFZ Single-Cell Sequencing Open Lab in designing and conducting the scRNA-Seq experiments. We wish to thank Alberto Diaz, Alicia Alonso, Maria Ramos and Kalman Somogyi for their suggestions on the manuscript. Schemes have been generated with BioRender.com. This work was supported by the Deutsches Zentrum für Lungenforschung (DZL, German Center for Lung Research # 82DZL004A4) to R.S and C.P.; Y.C and D.W. were supported by the Helmholtz Foundation. G.T.S. was supported by the Graduate College (Graduiertenkolleg, GRK) #2338 of the German Research Society (Deutsche Forschungsgemeinschaft, DFG), the target validation project for pharmaceutical development ALTERNATIVE of the German Ministry for Education and Research (Bundesministerium für Bildung und Forschung, BMBF), and a Translational Research Grant by the German Centre for Lung Research (DZL).

## Author Contributions

Y.C., G.T.S., S.C., C.P. and R.S., designed the experiments. Y.C. and S.C. performed the experiments and analysed the data. R.T. and J.H, conducted bioinformatic analyses, R.T., and P.L. developed bioinformatic methods. Y.C. and D.W. performed the TBWGBS. S.S. developed the method to detect mouse lung tumours by μCT. Y.C., S.C., R.T., C.P. and R.S. wrote the manuscript with comments from all authors.

## Conflicts of Interest

The authors declare no conflict of interest.

## STAR methods

### RESOURCE AVAILABILITY

#### Lead contact

Further information and requests for resources and reagents should be directed to and will be fulfilled by the lead contact, Rocio Sotillo.

#### Materials availability

### EXPERIMENTAL MODEL AND SUBJECT DETAILS

#### Mice, tamoxifen and adenoviral infection

All animal experiments were carried out following the guidelines of EU animal laws and approved by Baden-Wurttemberg, Germany (animal license No. G185-17, G265-19). *mT/mG* mice in C57BL/6 and FVB mixed background were kindly provided by Georgios Stathopoulos (University of Patras, Greece), *Hopx-CreERT2* line (ID: 017606) in 129 background and *Krt5-CreERT2* line in C57BL/6 and SJL mixed background (ID: 018394) were purchased from The Jackson Laboratory. *Sftpc-CreER-rtTA* was kindly provided by Harold Chapman (University of California, San Francisco, US), and *Scgb1a1-CreERT, Foxj1- CreERT* in C57BL/6 background were kindly provided by Brigit Hogan (Duke University, Durham, US). All animals were kept in the abovementioned background mixed with FVB.

To label epithelial cells in the lung, 6-12-week-old female and male mice were injected *i.p.* with 200 μg g^-1^ tamoxifen (Sigma, T5648, 30 mg ml^-1^ dissolved in corn oil) during 4 consecutive days. To select the optimal chasing period, we compared the number of labelled cells using 4 and 8 weeks chasing time in *Scgb1a1* mice. Since there were no differences of labelled cells in these two time points and previous data from the literature suggested that a period over 3 weeks was sufficient for TAM to label cells (Vaughan et al., 2015), we used 4 weeks of chasing time in all the experiments.

To induce *Eml4-Alk* rearrangement in the lung, mice were anaesthetized by intraperitoneal injection of 100 μg g^-1^ ketamine and 14 μg g^-1^ xylazine and intratracheally instilled with *Eml4-Alk* adenovirus, designed in the Ventura Lab (Maddalo et al., 2014) and purchased from Viraquest. The use of genetically modified organisms (GMO) was approved by the government of Baden-Wurttemberg, Germany (project No. 81078, 81155). Mice were randomly assigned to different experiments and investigators were not blind to the mice allocation during experiments and analysis.

Mice were killed by cervical dislocation and the lungs were perfused with 10 ml PBS through the right ventricle.

### METHOD DETAILS

#### Immunostaining

Mouse lungs were incubated with 10% formalin (Sigma, HT501128) on a tube rocker for 24 hours before further processing in a tissue processor (Leica ASP300S). Lungs were embedded in paraffin blocks and sectioned at 3μm thickness. For immunofluorescent staining, the following primary antibodies were used: ProSP-C (Millipore, AB3786, 1:500), CC10 (Santa Cruz, SC-9772, 1:500), acetylated tubulin (Sigma, T7451, 1:500), Cytokeratin 5 (Abcam, ab53121, 1:200), Podoplanin (Abcam, ab11936, 1:200), GFP (Cell Signalling Technology, 2956S, 1:200), GFP (Abcam, ab5450, 1:200), RFP (Rockland 600-401-379, 1:200), Aquaporin 5 (Abcam, ab78486, 1:1000) and Scgb3a2 (1:500, R&D systems, AF3465). Secondary antibodies were Alexa 488 donkey anti-rabbit IgG (Thermofisher, A21206, 1:500), Alexa 488 donkey anti-goat IgG (Abcam, ab150129, 1:500), Alexa 568 donkey anti-rabbit IgG (Thermofisher, A10042, 1:500), Alexa 568 donkey anti-goat IgG (Thermofisher, A11057, 1:500) and Alexa 568 donkey anti-mouse IgG (Abcam, ab175700, 1:500). Pictures were taken in a Leica SP5 confocal system and Tissuegnostic TissueFAX system. For immunohistochemical staining, the ABC kit (PK-6101) and DAB peroxidase substrate kit (SK-4100) from Vector Laboratories were used according to the manufacturer’s instructions. Primary antibodies were: GFP (Cell Signalling Technology, 2956S, 1:200), CCSP (Millipore, 07-623, 1:1,000), SPC (Millipore, AB3786, 1:500), STAT1 (Sigma, HPA000982, 1:500) and Lactotransferin (Sigma, 07-685, 1:500). Pictures were taken in a Zeiss Axioplan microscope and Tissuegnostic TissueFAX system.

#### Lung/tumour cell isolation and FACS sorting

For normal cells, lungs were minced into smaller pieces, and for tumour cells, visible nodules were carefully picked out, and the surrounding healthy tissue was removed. For TWGBS, both healthy lung and tumours were dissociated into single cells using the lung dissociation kit (Miltenyi Biotech, 130-095-927) in a gentleMACS Octo Dissociator (Miltenyi Biotech, 130-095-937). After depletion of red blood cells with the red blood cell lysis buffer (Sigma, R7757), cells were incubated with tumour cell isolation kit (Miltenyi Biotech, 130-110-187) following the manufacturer’s instruction and passed through the LS columns (Miltenyi Biotech, 130-042-401). Cells from the flow-through were collected and DAPI (1 μg ml^-1^) was added as a viability marker. Cells were then sorted in a BD FACSAria cell sorter and the DAPI;tdTomato; GFP^+^ population was collected for downstream applications.

For single-cell RNA sequencing, lungs were perfused with PBS, injected with 1 ml digestion cocktail (50 U ml^-1^ dispase, 250 U ml^-1^ collagenase, 5 U ml^-1^ elastase, 30 μg ml^-1^ DNAse I) through the trachea and cropped into small pieces. They were then incubated with 3ml of digestion cocktail on a tube rocker for 30 minutes under room temperature before being dissociated into small pieces by plungers in 10cm petri dishes. Samples were incubated 10min with 5ml DMEM with 10% FCS, 1% P/S and 100 μg ml^-1^ DNAse I at room temperature. For endpoint tumours, visible nodules were picked out after adjacent healthy tissue was removed. Tumour cells were processed into single cells using the lung dissociation kit and red blood cells lysis buffer as described above. Tumour cells were then incubated with CD31 and CD45 microbeads (Miltenyi Biotech, 130-110-187) following the manufacturer’s instructions and passed through the LS columns (Miltenyi Biotech, 130-042-401). Cells from the flow-through were incubated with DAPI (1 μg ml^-1^), CD45-PE (1 μg ml^-1^), CD31-PE (1 μg ml^-1^), EpCAM- BV711 (1 μg ml^-1^), CD24-BUV395 (1 μg ml^-1^), ß4-Alexa Fluor 647 (5 μg ml^-1^) and CD200- BV421 (1 μg ml^-1^) for further purification. Cells were then sorted in a BD FACSAria cell sorter and live EpCAM^+^; CD45^-^; CD31^-^; tdTomato^-^; GFP^+^; CD24^-^; ß4^+^; CD200^+^ population was collected for TAM, Cas9, 2wk-1, 2wk-2 and 4wk-1 groups and live EpCAM^+^; CD45^-^; CD31^-^; tdTomato^-^; GFP^+^ population was collected for 4wk-GFP, tumours. Collected cells were resuspended in 1,000 cells/μl before proceeding to library preparation.

#### DNA isolation and library preparation for bisulfite sequencing

Lung cell dissociation and sorting of GFP^+^ cells were done as mentioned above. Each sample was a pool of 1 to 4 mice. DNA was isolated with QIAamp DNA Micro Kit (Qiagen, 56304). Tagmentation-based whole genome bisulfite sequencing (TWGBS) libraries were generated as described previously (Wang et al., 2013) using 20-30 ng genomic DNA as input. Per sample, four sequencing libraries with different barcodes were prepared and pooled in equimolar amounts to a final concentration of 2-10 nM. Pools were sequenced paired-end, 125 bp, on one lane of a HiSeq2000 sequencer (Illumina). Raw fastq files were aligned to the mm10 reference genome using methylCtools (Hovestadt et al., 2014) as implemented by the Genomics and Proteomics Core Facility of the German Cancer Research Center.

#### DNA methylation analysis

Methylation levels were called by MethylDackel (https://github.com/dpryan79/MethylDackel). Due to the specifics of the TWGBS, the following parameters were used to remove M-bias: --nOB 2,11,11,2 --nOT 8,1,2,11. Quality control of the results included checking M-bias, bisulfite conversion rate and global methylation. Two samples were removed due to quality issues. Methylation data were analysed using R/Bioconductor 4.0 with packages Methrix (Mayakonda et al., 2020) and bsseq (Hansen et al., 2012). CpG sites overlapping with single nucleotide polymorphisms (SNPs) in any of the mouse strains were excluded based on data downloaded from Mouse Genome Project, database version 142 (Keane et al., 2011). The quality of the samples is shown in Supplementary Table 5.

For reference-free cell type deconvolution, we used MeDeCom, which allows the decomposition of DNA methylation into latent methylation components (Lutsik et al., 2017). Regions were selected based on the 15-state ChromHMM model for mouse lung, postnatal 0 days downloaded from ENCODE (Davis et al., 2018; ENCODE Project Consortium, 2012) (ENCSR538YJF). The 100000 most variable CpG sites overlapping with poised or bivalent enhancers were included in the model. MeDeCom model was run using multiple lambda and K parameters with the following arguments: NINIT = 10, NFOLDS = 10, ITERMAX = 300. The final model with K=4 and lambda=0.0001 was selected based on cross-validation error. Labelled tumours were re-categorized based on their suspected cell-of-origin. Tumours with >10% proportion in LMC1 or LMC3 were categorized as Club or AT2 originating, respectively. To identify gene promoters negatively associated with the LMCs, we calculated the Pearson correlation coefficient between LMC proportions and promoter methylation. Using this as ranking, we performed a Gene Set Enrichment Analysis (GSEA), as implemented in the fgsea package (Korotkevich et al., 2021) with the following parameters: minSize = 3, maxSize = 500, nperm=1000. For each set of marker genes, the running score was visualized.

Differential methylation calling was performed with the DSS package (Park and Wu, 2016). The data was smoothed with a smoothing span of 500. Dispersion of the groups was assumed non-equal. Regions were assigned as differentially methylated based on the following parameters: delta=0, p.threshold=0.001, minlen=50, minCG=3, dis.merge=50, pct.sig=0.5. DMRs were annotated using ChromHMM 15-state model for mouse lung, postnatal 0 days (ENCSR538YJF), using annotatr package (Cavalcante and Sartor, 2017). Visualizations were created using ggplot2 (Wickham), ComplexHeatmap (Gu et al., 2016) and Gviz (Hahne and Ivanek, 2016) packages.

Transcription factor binding motif enrichment analysis was performed with Homer (Heinz et al., 2010) based on DMRs with the following parameters: -len 8,10,12 -size 100 -S 8 -cache 6921 -fdr 0.

#### Single-cell RNA sequencing and data analysis

The time points and gating strategy used to collect the samples are shown in Figure 5A and Supplementary Figure 5A. Lung cells were dissociated as mentioned above and GFP^+^ cells were harvested as indicated in Supplementary Figure 5A. Libraries were prepared using Chromium Next GEM Single Cell 3’ GEM, Library & Gel Bead Kit v3.1 (10x Genomics) according to manufacturer’s instruction. Gene expression counts were acquired using Cell Ranger 3.1 count from 10x Genomics with its default settings. The gene expression counts were analysed in R 4.0.3 using Seurat 3.2.2 (Stuart et al., 2019) unless indicated otherwise. Based on visual inspection, the following number of detected gene thresholds were used to exclude bad quality cells among the samples: 1000-6000 for TAM, 1000-6000 for Cas9, 1000-6000 for 2 wk-1, 2000-6000 for 2 wk-2, 1500-6000 for 4 wk-1, 2000-6000 for 4-wk GFP^+^, 1000-6000 for tumour. Cells with more than 10% mitochondrial genes were removed from all samples. Samples were normalized using a scaling factor of 10000. Variable features were selected using the FindVariableFeatures function of Seurat.

The cell-cycle stage of the cells was calculated using the CellCycleScoring function of Seurat with a publicly available dataset (Macosko et al., 2015). Genes were converted from human to mouse using the BioMart database with biomaRt R package (Durinck et al., 2005; 2009).

Batch effect correction was done using Harmony (Korsunsky et al., 2019) based on the PCA dimensionality reduction. Based on the resulted embeddings, we performed Uniform Manifold Approximation and Projection (UMAP) (McInnes et al., 2018) and clustering. Clusters were calculated using a shared nearest neighbour (SNN) modularity optimization-based clustering algorithm. First, the k-nearest neighbours were calculated based on the first 30 dimensions of the Harmony embedding and an SNN graph was constructed. An optimal number of clusters was selected based on a suggestion by an elbow plot, as implemented in Seurat. Clustering was performed using the Louvain algorithm with a resolution of 0.6. Overexpressed markers of the clusters were selected if they were expressed in more than 25% of the cluster with a log fold change (FC) of 0.25.

After the initial analysis, a small cluster of Ciliated cells marked by high expression of *Foxj1* was identified and excluded. The above process was then repeated without Ciliated cells.

*Alk* expression was calculated by counting the reads overlapping with the part of *Alk* gene affected by the translocation (chr17: 71867045-71898183). The counts were lognormalized to the number of transcripts in each cell.

Consensus non-negative matrix factorization (cNMF) (Kotliar et al., 2019) v.1.2 was run using 100 iterations and the 2000 most variable genes. Based on the optimization, a model with 9 components was selected. For the consensus estimates of the programs a local density threshold of 0.01 was used. The other parameters were used as default.

Overrepresentation analysis of the genes representing the modules based on KEGG (Kanehisa et al., 2021) and WikiPathways (Martens et al., 2021) databases were performed using the clusterProfiler Bioconductor package (Yu et al., 2012).

Module and signature scores were calculated using the AddModuleScore function of Seurat based on the default parameters.

RNA velocity models the direction and speed of individual cells in the gene expression space by estimating the ratio of spliced and unspliced mRNA. Using this ratio, it predicts the future state of individual cells on a short timescale (La Manno et al., 2018). Here, we calculated the velocity using scVelo 0.2.3 (Bergen et al., 2020). Since our data was batch corrected using Harmony the calculated velocity was projected on the UMAP based on the Harmony embedding.

Partition-based graph abstraction (PAGA) (Wolf et al., 2019) generates a topology-reserving map of single cells. Here we used PAGA as implemented in scanpy 1.7.2 (Wolf et al., 2018). For the standalone PAGA visualization, we used the algorithm on the Harmony embedding. We also used RNA velocity to direct the PAGA edges, as implemented in scVelo (Bergen et al., 2020; Wolf et al., 2019).

#### μCT imaging

μCT imaging was performed using the Inveon multi-modality μPET/SPECT/CT system (Siemens Medical Solutions, Knoxville, USA). Acquisitions covering the thorax and lungs were performed using a tube voltage of 80 kV and a tube current of 500 μA. A total of 360 projections were acquired over 360° with an integration time of 200 ms each. The detector was operated using a 4 x 4 binning mode resulting in a resolution of approximately 100 μm in the centre of rotation. Image reconstruction with isotropic resolution was performed using the Feldkamp algorithm with a Shepp-Logan kernel onto a 512 x 512 x 928 grid with appropriately sized voxels. Image analysis was performed using ImageJ.

#### Quantification and statistical analysis

The statistical information for the experiments are detailed in the text, figure legends, and figures.

Correlation between LMCs and gene promoter methylation was calculated using Pearson correlation.

If not otherwise stated, p value <0.05 was considered significant. False discovery rate (FDR) was used as a multiple test correction method, where appropriate. In this case, FDR q < 0.05 was considered significant.

## Supplemental Figure legends

**Supplementary Figure 1.**
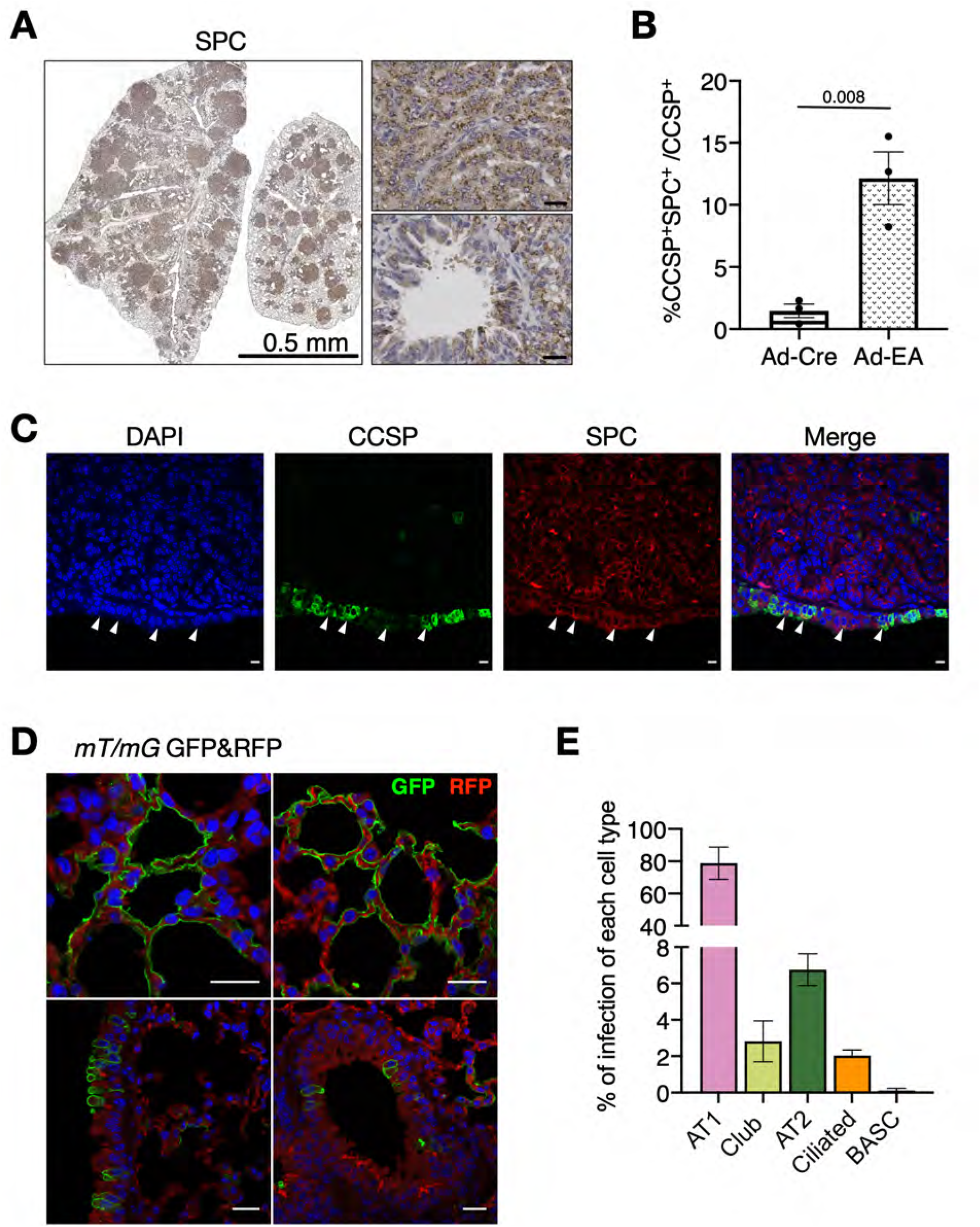
A) SPC staining of a representative *Eml4-Alk* tumour bearing lung showing that all tumours are positive for SPC. Scale bars 0.5mm and 100μm. B) Percentage of CCSP^+^SPC^+^ double-positive cells in the CCSP^+^ population in control mice (n=3, Ad-Cre) and in mice with *Eml4-Alk* induced tumours (n=3, Ad-EA). Unpaired t test with Welch’s correction is used to calculate the p value. C) Double-positive CCSP^+^SPC^+^ cells are found in the bronchioles during tumorigenesis. Arrowheads show double-positive cells. Antibodies are indicated. Scale bars: 10μm. D) GFP and RFP immunofluorescent staining on lung sections from *mT/mG* mice after infection with Ad-Cre (alveolar region, upper images; bronchiolar region, bottom images). Scale bars 20μm. E) Percentage of each cell type infected after Ad-Cre installation in *mT/mG* mice.

**Supplementary Figure 2.**
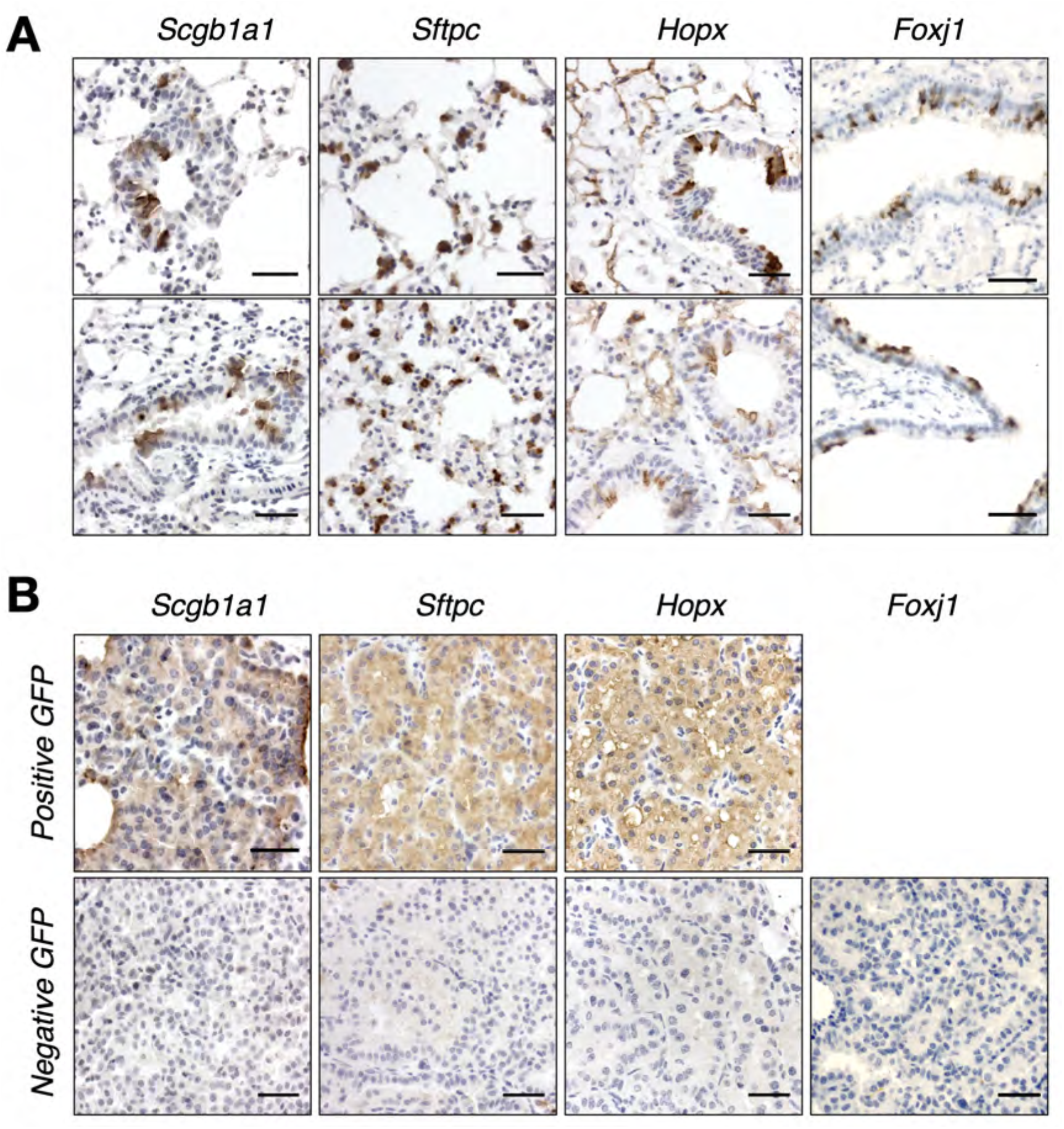
A) GFP immunohistochemical staining on lung sections from lineage-tracing mice as indicated. Scale bar 100μm. B) GFP immunohistochemical staining on lung tumours originated in the indicated lineage-tracing mice showing positive tumours in the upper panel and negative tumours in the lower panel. Scale bar 100μm.

**Supplementary Figure 3.**
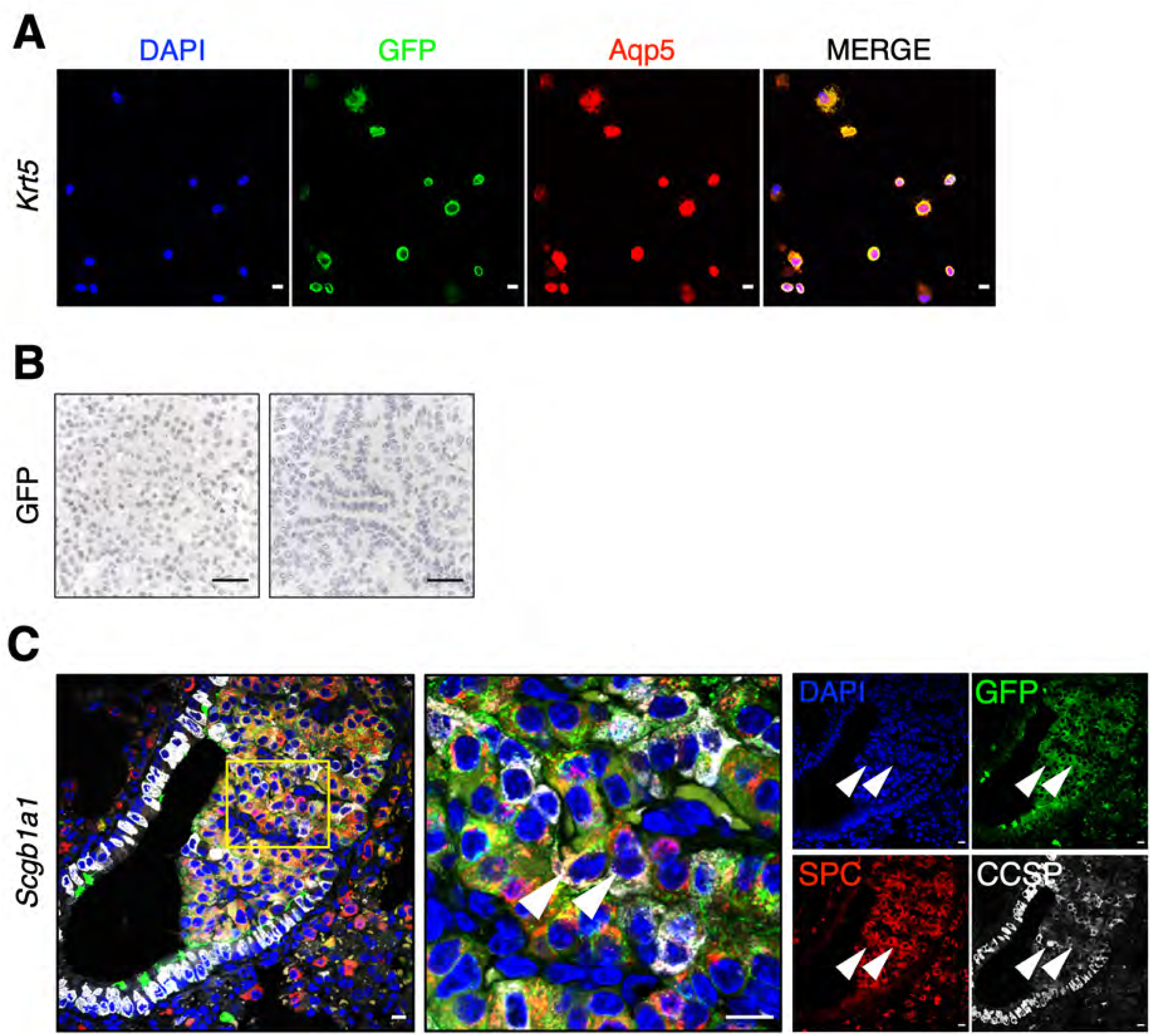
A) GFP and Aquaporin 5 (AQP5) immunofluorescent staining on FACS-sorted GFP^+^ cells from *Krt5* mice. Overly and single-channel images are sequentially presented. Scale bars: 5μm. B) GFP immunohistochemical staining on the tumours from *Krt5* mice. Scale bars: 50μm. C) GFP, SPC and CCSP immunofluorescent staining of early lesions originated in *Scgb1a1* lineage-tracing mice. Magnification of the highlighted area is shown on the right and arrowheads indicate Club cells under lineage switch. Scale bar: 10μm.

**Supplementary Figure 4.**
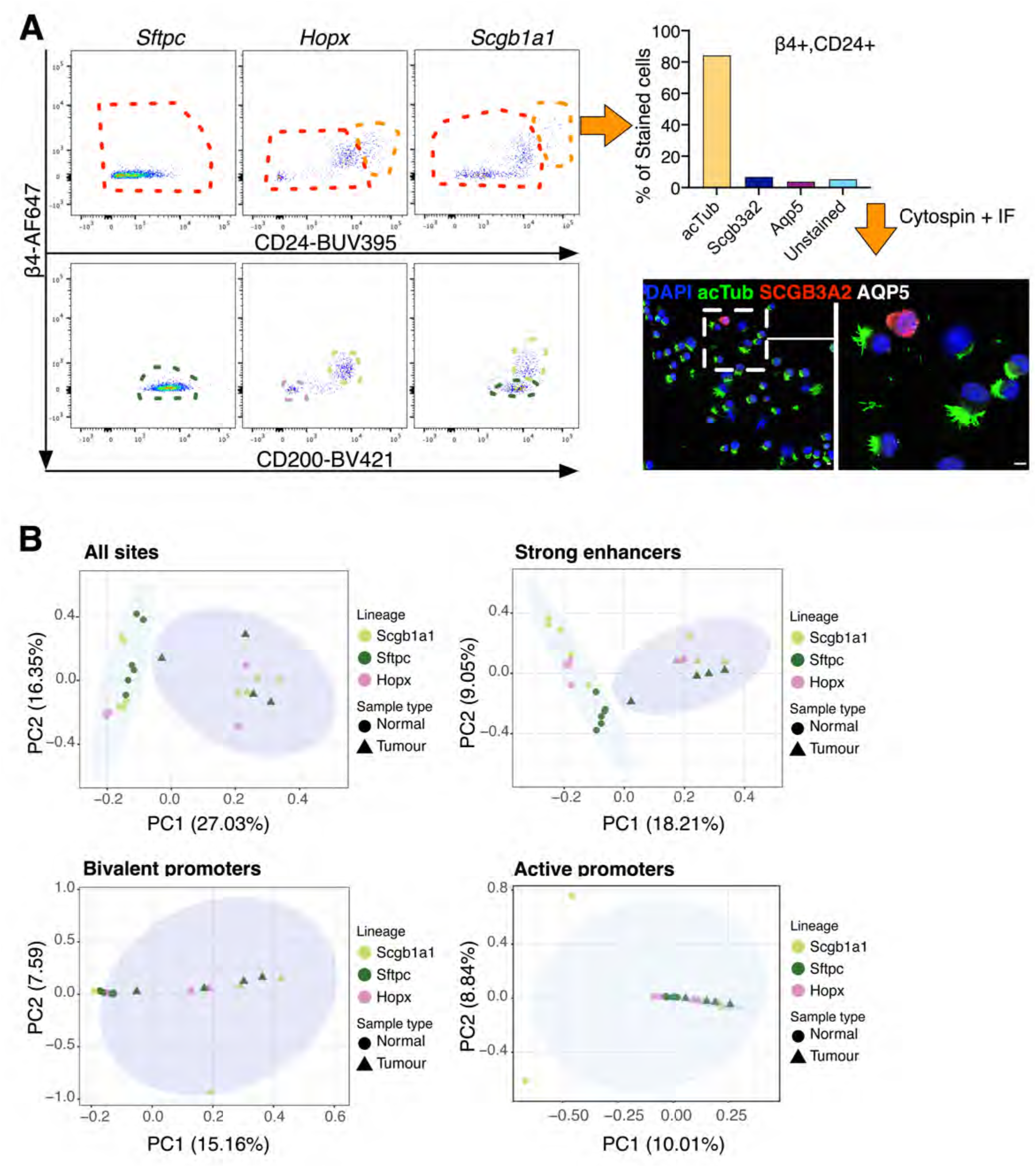
A) Sorting strategy of the isolated GFP^+^ cells in the indicated lineage-tracing models for tagmentation-based whole-genome bisulfite sequencing (TWGBS). Dashed lines indicate different cell types that compose the sorted GFP^+^ population: orange, Ciliated cells (β4^+^, CD24^+^); red, non-ciliated cells (β4^+^, CD24^-^). Out of the non-ciliated cells (red gate), there are AT2 cells (dark green; β4^-^, CD200^-^), Club cells (light green; β4^+^, CD200^+^), and AT1 cells (pink; β4^-^, CD200^-^). Further demonstration of the Ciliated cell population is provided through immunofluorescent staining of the sorted β4^+^, CD24^+^ population using the indicated cell type specific markers (acTub for Ciliated cells, SCGB3A2 for Club cells and AQP5 for AT1 cells). A total of 266 cells from 3 views was used to generate the bar plot. B) Principal component analysis plot of DNA methylation. CpG sites were selected in different ways: top variable sites from all regions (all sites), as well as sites overlapping with strong enhancers (En-Sd, En-Sp), bivalent promoters (Pr-B), or active promoters (Pr-A). The functional regions were defined based on ChromHMM tracks from postnatal 0-day old mouse lung (ENCODE, ENCSR538YJF).

**Supplementary Figure 5.**
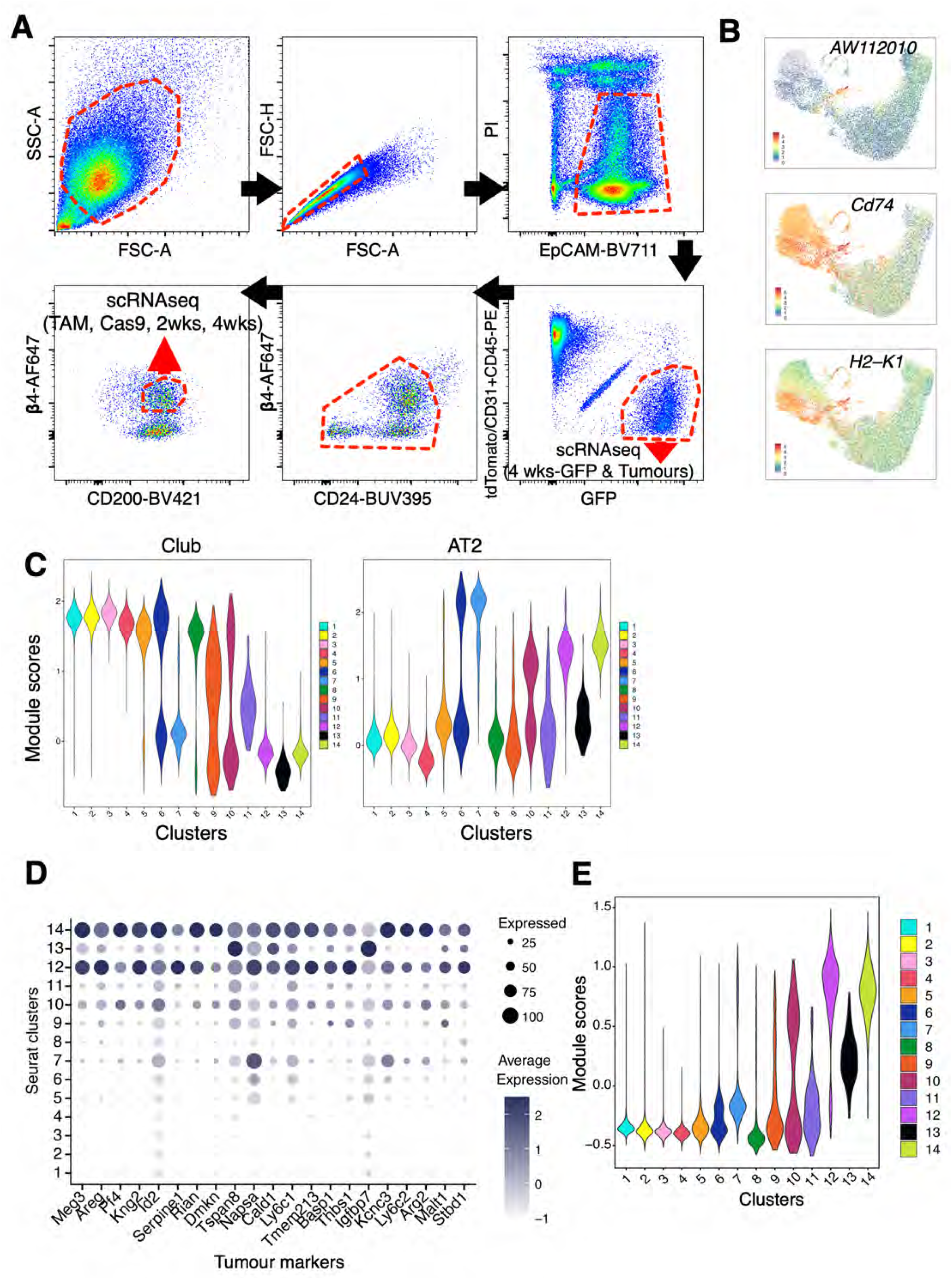
A) Sorting strategy to obtain GFP labelled Club cells in the *Scgb1a1* line for scRNA-seq. Dashed lines indicate the gates applied to select the subpopulations from the parental cells. Arrows indicate the cell populations that were sequenced. B) UMAP embedding with cells coloured by the expression levels of *AW112010, Cd74,* and *H2-K1.* X- and y-axes represent the first and second UMAP dimensions (UMAP_1 and UMAP_2), respectively. C) Violin plots showing the expression module score of Club (left) and AT2 (right) signatures by the identified clusters. The signatures were used as described by Strunz et al. (Strunz et al., 2020). D) Dot plot showing the high expression of tumour genes mainly in clusters 12 and 14. The size of the dots represents the percentage of cells in the cluster that expresses the gene, while the intensity of the colour shows the average expression of the gene. E) Violin plot indicating the module score of the tumour signature by the identified clusters (signature composed by the genes in C).

**Supplementary Figure 6.**
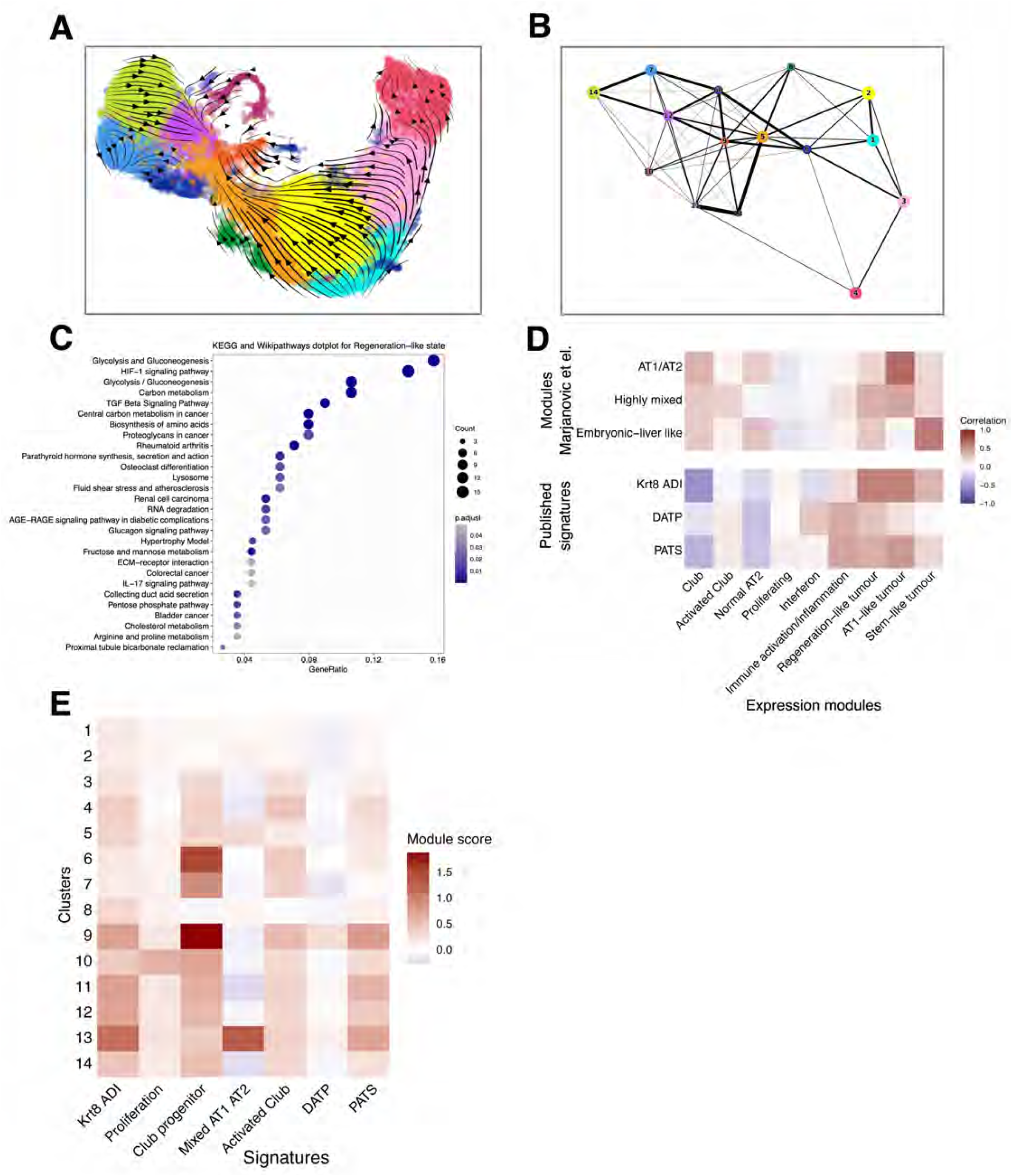
A) RNA velocity plot projected to the UMAP embedding indicating the directional progression of transcriptional states of the clusters. B) Partition-based graph abstraction (PAGA) analysis projected to the previously identified clusters. Cluster nodes are connected with weighted edges (width of the connections), with the weights representing the connectivity between the clusters. C) Pathway enrichment analysis based on KEGG and WikiPathways for the regeneration-like tumour activity module, identified by cNMF. The size of the dots shows the number of genes among the top 200 genes in the activity module. The colour represents the level of significance. The x-axis depicts the gene ratio between the number of genes in the pathway and the number of genes among the selected ones. D) Heatmap of the correlation between the identified activity programs and the programs identified by Marjanovic et al. or previously published signatures (Choi et al., 2020; Kobayashi et al., 2020; Strunz et al., 2020) (Supplementary Table 4). In the upper part of the figure, the colours represent the correlation between the z-score unit gene expression of the gene activity programs calculated using cNMF, while the lower part shows the correlation of module scores calculated using the top 10 genes of the activity program or the genes of the published signatures. E) Module scores of previously identified signatures by cluster (Choi et al., 2020; Kobayashi et al., 2020; Strunz et al., 2020).

## Notes

### Competing Interest Statement

The authors have declared no competing interest.

## References

Alonso-Curbelo, D., Ho, Y.-J., Burdziak, C., Maag, J.L.V., Morris, J.P., Chandwani, R., Chen, H.-A., Tsanov, K.M., Barriga, F.M., Luan, W., et al. (2021). A gene-environment-induced epigenetic program initiates tumorigenesis. Nature 17, 594–597.

Angelidis, I., Simon, L.M., Fernandez, I.E., Strunz, M., Mayr, C.H., Greiffo, F.R., Tsitsiridis, G., Ansari, M., Graf, E., Strom, T.-M., et al. (2019). An atlas of the aging lung mapped by single cell transcriptomics and deep tissue proteomics. Nat Commun 10, 963–17.

Bergen, V., Lange, M., Peidli, S., Wolf, F.A., and Theis, F.J. (2020). Generalizing RNA velocity to transient cell states through dynamical modeling. Nat. Biotechnol. 38, 1408–1414.

Capper, D., Jones, D.T.W., Sill, M., Hovestadt, V., Schrimpf, D., Sturm, D., Koelsche, C., Sahm, F., Chavez, L., Reuss, D.E., et al. (2018). DNA methylation-based classification of central nervous system tumours. Nature 555, 469–474.

Cavalcante, R.G., and Sartor, M.A. (2017). annotatr: genomic regions in context. Bioinformatics 33, 2381–2383.

Chapman, H.A., Li, X., Alexander, J.P., Brumwell, A., Lorizio, W., Tan, K., Sonnenberg, A., Wei, Y., and Vu, T.H. (2011). Integrin α6β4 identifies an adult distal lung epithelial population with regenerative potential in mice. J. Clin. Invest. 121, 2855–2862.

Choi, J., Park, J.-E., Tsagkogeorga, G., Yanagita, M., Koo, B.-K., Han, N., and Lee, J.-H. (2020). Inflammatory Signals Induce AT2 Cell-Derived Damage-Associated Transient Progenitors that Mediate Alveolar Regeneration. Cell Stem Cell 27, 366–382.e367.

Davis, C.A., Hitz, B.C., Sloan, C.A., Chan, E.T., Davidson, J.M., Gabdank, I., Hilton, J.A., Jain, K., Baymuradov, U.K., Narayanan, A.K., et al. (2018). The Encyclopedia of DNA elements (ENCODE): data portal update. Nucleic Acids Research 46, D794–D801.

Desai, T.J., Brownfield, D.G., and Krasnow, M.A. (2014). Alveolar progenitor and stem cells in lung development, renewal and cancer. Nature 507, 190–194.

Dor, Y., and Cedar, H. (2018). Principles of DNA methylation and their implications for biology and medicine. Lancet 392, 777–786.

Dost, A.F.M., Moye, A.L., Vedaie, M., Tran, L.M., Fung, E., Heinze, D., Villacorta-Martin, C., Huang, J., Hekman, R., Kwan, J.H., et al. (2020). Organoids Model Transcriptional Hallmarks of Oncogenic KRAS Activation in Lung Epithelial Progenitor Cells. Cell Stem Cell.

Durinck, S., Moreau, Y., Kasprzyk, A., Davis, S., De Moor, B., Brazma, A., and Huber, W. (2005). BioMart and Bioconductor: a powerful link between biological databases and microarray data analysis. Bioinformatics 21, 3439–3440.

Durinck, S., Spellman, P.T., Birney, E., and Huber, W. (2009). Mapping identifiers for the integration of genomic datasets with the R/Bioconductor package biomaRt. Nature Protocols 4, 1184–1191.

ENCODE Project Consortium (2012). An integrated encyclopedia of DNA elements in the human genome. Nature 489, 57–74.

Ferone, G., Lee, M.C., Sage, J., and Berns, A. (2020). Cells of origin of lung cancers: lessons from mouse studies. Genes & Development 34, 1017–1032.

Gu, Z., Eils, R., and Schlesner, M. (2016). Complex heatmaps reveal patterns and correlations in multidimensional genomic data. Bioinformatics 32, 2847–2849.

Hahne, F., and Ivanek, R. (2016). Visualizing Genomic Data Using Gviz and Bioconductor. In Statistical Genomics, (Humana Press, New York, NY), pp. 335–351.

Han, X., Wang, R., Zhou, Y., Fei, L., Sun, H., Lai, S., Saadatpour, A., Zhou, Z., Chen, H., Ye, F., et al. (2018). Mapping the Mouse Cell Atlas by Microwell-Seq. Cell 172, 1091–1107.e17.

Hansen, K.D., Langmead, B., and Irizarry, R.A. (2012). BSmooth: from whole genome bisulfite sequencing reads to differentially methylated regions. Genome Biol 13, R83–10.

Heinz, S., Benner, C., Spann, N., Bertolino, E., Lin, Y.C., Laslo, P., Cheng, J.X., Murre, C., Singh, H., and Glass, C.K. (2010). Simple combinations of lineage-determining transcription factors prime cis-regulatory elements required for macrophage and B cell identities. Molecular Cell 38, 576–589.

Hovestadt, V., Jones, D.T.W., Picelli, S., Wang, W., Kool, M., Northcott, P.A., Sultan, M., Stachurski, K., Ryzhova, M., Warnatz, H.-J., et al. (2014). Decoding the regulatory landscape of medulloblastoma using DNA methylation sequencing. Nature 510, 537–541.

Indra, A.K., Warot, X., Brocard, J., Bornert, J.-M., Xiao, J.H., Chambon, P., and Metzger, D. (1999). Temporally-controlled site-specific mutagenesis in the basal layer of the epidermis: comparison of the recombinase activity of the tamoxifen-inducible Cre-ER(T) and Cre-ER(T2) recombinases. Nucleic Acids Research 27, 4324–4327.

Jain, R., Barkauskas, C.E., Takeda, N., Bowie, E.J., Aghajanian, H., Wang, Q., Padmanabhan, A., Manderfield, L.J., Gupta, M., Li, D., et al. (2015). Plasticity of Hopx(+) type I alveolar cells to regenerate type II cells in the lung. Nat Commun 6, 6727.

Kanehisa, M., Furumichi, M., Sato, Y., Ishiguro-Watanabe, M., and Tanabe, M. (2021). KEGG: integrating viruses and cellular organisms. Nucleic Acids Research 49, D545–D551.

Karamouzis, M.V., Konstantinopoulos, P.A., and Papavassiliou, A.G. (2007). The activator protein-1 transcription factor in respiratory epithelium carcinogenesis. Mol. Cancer Res. 5, 109–120.

Kathiriya, J.J., Brumwell, A.N., Jackson, J.R., Tang, X., and Chapman, H.A. (2020). Distinct Airway Epithelial Stem Cells Hide among Club Cells but Mobilize to Promote Alveolar Regeneration. Cell Stem Cell 26, 346–358.e4.

Keane, T.M., Goodstadt, L., Danecek, P., White, M.A., Wong, K., Yalcin, B., Heger, A., Agam, A., Slater, G., Goodson, M., et al. (2011). Mouse genomic variation and its effect on phenotypes and gene regulation. Nature 477, 289–294.

Kim, C.F.B., Jackson, E.L., Woolfenden, A.E., Lawrence, S., Babar, I., Vogel, S., Crowley, D., Bronson, R.T., and Jacks, T. (2005). Identification of bronchioalveolar stem cells in normal lung and lung cancer. Cell 121, 823–835.

Kim, M., and Costello, J. (2017). DNA methylation: An epigenetic mark of cellular memory. Exp Mol Med 49, e322–e322.

Kobayashi, Y., Tata, A., Konkimalla, A., Katsura, H., Lee, R.F., Ou, J., Banovich, N.E., Kropski, J.A., and Tata, P.R. (2020). Persistence of a regeneration-associated, transitional alveolar epithelial cell state in pulmonary fibrosis. Nat. Cell Biol. 22, 934–946.

Korotkevich, G., Sukhov, V., Budin, N., Shpak, B., Artyomov, M.N., and Sergushichev, A. (2021). Fast gene set enrichment analysis. bioRxiv 060012.

Korsunsky, I., Millard, N., Fan, J., Slowikowski, K., Zhang, F., Wei, K., Baglaenko, Y., Brenner, M., Loh, P.-R., and Raychaudhuri, S. (2019). Fast, sensitive and accurate integration of single-cell data with Harmony. Nat Meth 16, 1289–1296.

Kotliar, D., Veres, A., Nagy, M.A., Tabrizi, S., Hodis, E., Melton, D.A., and Sabeti, P.C. (2019). Identifying gene expression programs of cell-type identity and cellular activity with single-cell RNA-Seq. eLife 8.

La Manno, G., Soldatov, R., Zeisel, A., Braun, E., Hochgerner, H., Petukhov, V., Lidschreiber, K., Kastriti, M.E., Lönnerberg, P., Furlan, A., et al. (2018). RNA velocity of single cells. Nature 560, 494–498.

LaFave, L.M., Kartha, V.K., Ma, S., Meli, K., Del Priore, I., Lareau, C., Naranjo, S., Westcott, P.M.K., Duarte, F.M., Sankar, V., et al. (2020). Epigenomic State Transitions Characterize Tumor Progression in Mouse Lung Adenocarcinoma. Cancer Cell 1–45.

Li, S., Wang, Y., Zhang, Y., Lu, M.M., DeMayo, F.J., Dekker, J.D., Tucker, P.W., and Morrisey, E.E. (2012). Foxp1/4 control epithelial cell fate during lung development and regeneration through regulation of anterior gradient 2. Development 139, 2500–2509.

Lipka, D.B., Witte, T., Toth, R., Yang, J., Wiesenfarth, M., Nöllke, P., Fischer, A., Brocks, D., Gu, Z., Park, J., et al. (2017). RAS-pathway mutation patterns define epigenetic subclasses in juvenile myelomonocytic leukemia. Nat Commun 8, 2126–14.

Liu, Q., Liu, K., Cui, G., Huang, X., Yao, S., Guo, W., Qin, Z., Li, Y., Yang, R., Pu, W., et al. (2019). Lung regeneration by multipotent stem cells residing at the bronchioalveolar-duct junction. Nat. Genet. 51, 728–738.

Lutsik, P., Slawski, M., Gasparoni, G., Vedeneev, N., Hein, M., and Walter, J. (2017). MeDeCom: discovery and quantification of latent components of heterogeneous methylomes. Genome Biol 18, 55–20.

Macosko, E.Z., Basu, A., Satija, R., Nemesh, J., Shekhar, K., Goldman, M., Tirosh, I., Bialas, A.R., Kamitaki, N., Martersteck, E.M., et al. (2015). Highly Parallel Genome-wide Expression Profiling of Individual Cells Using Nanoliter Droplets. Cell 161, 1202–1214.

Maddalo, D., Manchado, E., Concepcion, C.P., Bonetti, C., Vidigal, J.A., Han, Y.-C., Ogrodowski, P., Crippa, A., Rekhtman, N., de Stanchina, E., et al. (2014). In vivo engineering of oncogenic chromosomal rearrangements with the CRISPR/Cas9 system. Nature 516, 423–427.

Mainardi, S., Mijimolle, N., Francoz, S., Vicente-Dueñas, C., Sánchez-García, I., and Barbacid, M. (2014). Identification of cancer initiating cells in K-Ras driven lung adenocarcinoma. Proc. Natl. Acad. Sci. U.S.a. 111, 255–260.

Marjanovic, N.D., Hofree, M., Chan, J.E., Canner, D., Wu, K., Trakala, M., Hartmann, G.G., Smith, O.C., Kim, J.Y., Evans, K.V., et al. (2020). Emergence of a High-Plasticity Cell State during Lung Cancer Evolution. Cancer Cell 38, 229–246.e13.

Martens, M., Ammar, A., Riutta, A., Waagmeester, A., Slenter, D.N., Hanspers, K., A Miller, R., Digles, D., Lopes, E.N., Ehrhart, F., et al. (2021). WikiPathways: connecting communities. Nucleic Acids Research 49, D613–D621.

Mayakonda, A., Schönung, M., Hey, J., Batra, R.N., Feuerstein-Akgoz, C., Köhler, K., Lipka, D.B., Sotillo, R., Plass, C., Lutsik, P., et al. (2020). Methrix: an R/bioconductor package for systematic aggregation and analysis of bisulfite sequencing data. Bioinformatics.

McInnes, L., Healy, J., and Melville, J. (2018). UMAP: Uniform Manifold Approximation and Projection for Dimension Reduction.

Montoro, D.T., Haber, A.L., Biton, M., Vinarsky, V., Lin, B., Birket, S.E., Yuan, F., Chen, S., Leung, H.M., Villoria, J., et al. (2018). A revised airway epithelial hierarchy includes CFTR-expressing ionocytes. Nature 560, 319–324.

Muzumdar, M.D., Tasic, B., Miyamichi, K., Li, L., and Luo, L. (2007). A global double-fluorescent Cre reporter mouse. Genesis 45, 593–605.

Oakes, C.C., Seifert, M., Assenov, Y., Gu, L., Przekopowitz, M., Ruppert, A.S., Wang, Q., Imbusch, C.D., Serva, A., Koser, S.D., et al. (2016). DNA methylation dynamics during B cell maturation underlie a continuum of disease phenotypes in chronic lymphocytic leukemia. Nat. Genet. 48, 253–264.

Paranjapye, A., Mutolo, M.J., Ebron, J.S., Leir, S.-H., and Harris, A. (2020). The FOXA1 transcriptional network coordinates key functions of primary human airway epithelial cells. Am J Physiol Lung Cell Mol Physiol 319, L126–L136.

Park, Y., and Wu, H. (2016). Differential methylation analysis for BS-seq data under general experimental design. Bioinformatics 32, 1446–1453.

Plasschaert, L.W., Žilionis, R., Choo-Wing, R., Savova, V., Knehr, J., Roma, G., Klein, A.M., and Jaffe, A.B. (2018). A single-cell atlas of the airway epithelium reveals the CFTR-rich pulmonary ionocyte. Nature 560, 377–381.

Rawlins, E.L., Okubo, T., Xue, Y., Brass, D.M., Auten, R.L., Hasegawa, H., Wang, F., and Hogan, B.L.M. (2009). The role of Scgb1a1+ Clara cells in the long-term maintenance and repair of lung airway, but not alveolar, epithelium. Cell Stem Cell 4, 525–534.

Rawlins, E.L., Ostrowski, L.E., Randell, S.H., and Hogan, B.L.M. (2007). Lung development and repair: contribution of the ciliated lineage. Proceedings of the National Academy of Sciences 104, 410–417.

Rock, J.R., Onaitis, M.W., Rawlins, E.L., Lu, Y., Clark, C.P., Xue, Y., Randell, S.H., and Hogan, B.L.M. (2009). Basal cells as stem cells of the mouse trachea and human airway epithelium. Proceedings of the National Academy of Sciences 106, 12771–12775.

Rowbotham, S.P., and Kim, C.F. (2014). Diverse cells at the origin of lung adenocarcinoma. Proc. Natl. Acad. Sci. U.S.a. 111, 4745–4746.

Salwig, I., Spitznagel, B., Vazquez Armendariz, A.I., Khalooghi, K., Guenther, S., Herold, S., Szibor, M., and Braun, T. (2019). Bronchioalveolar stem cells are a main source for regeneration of distal lung epithelia in vivo. Embo J. 38.

Scherer, M., Nazarov, P.V., Toth, R., Sahay, S., Kaoma, T., Maurer, V., Plass, C., Lengauer, T., Walter, J., and Lutsik, P. (2020). Reference-free deconvolution of complex DNA methylation data - a systematic protocol. bioRxiv 9, 1341–26.

Sill, M., Plass, C., Pfister, S.M., and Lipka, D.B. (2020). Molecular tumor classification using DNA methylome analysis. Human Molecular Genetics 29, R205–R213.

Spella, M., Lilis, I., Pepe, M.A.A., Chen, Y., Armaka, M., Lamort, A.-S., Zazara, D.E., Roumelioti, F., Vreka, M., Kanellakis, N.I., et al. (2019). Club cells form lung adenocarcinomas and maintain the alveoli of adult mice. eLife 8.

Strunz, M., Simon, L.M., Ansari, M., Kathiriya, J.J., Angelidis, I., Mayr, C.H., Tsidiridis, G., Lange, M., Mattner, L.F., Yee, M., et al. (2020). Alveolar regeneration through a Krt8+ transitional stem cell state that persists in human lung fibrosis. Nat Commun 11, 3559–20.

Stuart, T., Butler, A., Hoffman, P., Hafemeister, C., Papalexi, E., Mauck, W.M., Hao, Y., Stoeckius, M., Smibert, P., and Satija, R. (2019). Comprehensive Integration of Single-Cell Data. Cell 177, 1888–1902.e21.

Sutherland, K.D., Song, J.-Y., Kwon, M.C., Proost, N., Zevenhoven, J., and Berns, A. (2014). Multiple cells-of-origin of mutant K-Ras-induced mouse lung adenocarcinoma. Proceedings of the National Academy of Sciences 111, 4952–4957.

Takeda, N., Jain, R., Leboeuf, M.R., Wang, Q., Lu, M.M., and Epstein, J.A. (2011). Interconversion between intestinal stem cell populations in distinct niches. Science 334, 1420–1424.

Tata, P.R., Mou, H., Pardo-Saganta, A., Zhao, R., Prabhu, M., Law, B.M., Vinarsky, V., Cho, J.L., Breton, S., Sahay, A., et al. (2013). Dedifferentiation of committed epithelial cells into stem cells in vivo. Nature 503, 218–223.

Touzart, A., Mayakonda, A., Smith, C., Hey, J., Toth, R., Cieslak, A., Andrieu, G.P., Tran Quang, C., Latiri, M., Ghysdael, J., et al. (2021). Epigenetic analysis of patients with T-ALL identifies poor outcomes and a hypomethylating agent-responsive subgroup. Science Translational Medicine 13.

Travaglini, K.J., Nabhan, A.N., Penland, L., Sinha, R., Gillich, A., Sit, R.V., Chang, S., Conley, S.D., Mori, Y., Seita, J., et al. (2019). A molecular cell atlas of the human lung from single cell RNA sequencing. bioRxiv 1–76.

Vaughan, A.E., Brumwell, A.N., Xi, Y., Gotts, J.E., Brownfield, D.G., Treutlein, B., Tan, K., Tan, V., Liu, F.C., Looney, M.R., et al. (2015). Lineage-negative progenitors mobilize to regenerate lung epithelium after major injury. Nature 517, 621–625.

Wang, Q., Gu, L., Adey, A., Radlwimmer, B., Wang, W., Hovestadt, V., Bähr, M., Wolf, S., Shendure, J., Eils, R., et al. (2013). Tagmentation-based whole-genome bisulfite sequencing. Nature Protocols 8, 2022–2032.

Wickham, H. ggplot2: Elegant Graphics for Data Analysis.

Wierzbinska, J.A., Toth, R., Ishaque, N., Rippe, K., Mallm, J.-P., Klett, L.C., Mertens, D., Zenz, T., Hielscher, T., Seifert, M., et al. (2020). Methylome-based cell-of-origin modeling (Methyl-COOM) identifies aberrant expression of immune regulatory molecules in CLL. Genome Med 12, 29–19.

Wolf, F.A., Angerer, P., and Theis, F.J. (2018). SCANPY: large-scale single-cell gene expression data analysis. Genome Biol 19, 15–15.

Wolf, F.A., Hamey, F.K., Plass, M., Solana, J., Dahlin, J.S., Göttgens, B., Rajewsky, N., Simon, L., and Theis, F.J. (2019). PAGA: graph abstraction reconciles clustering with trajectory inference through a topology preserving map of single cells. Genome Biol 20, 59–59.

Xu, X., Rock, J.R., Lu, Y., the, C.F.P.O., 2012 (2012). Evidence for type II cells as cells of origin of K-Ras-induced distal lung adenocarcinoma. National Acad Sciences, 109, 4910–4915.

Yau, C., Wolf, D.M., Lazar, A.J., Drill, E., Shen, R., Taylor, A.M., Thorsson, V., Wiznerowicz, M., Robertson, A.G., Schneider, B.G., et al. (2018). Cell-of-Origin Patterns Dominate the Molecular Classification of 10,000 Tumors from 33 Types of Cancer. Cell 173, 291–304.e296.

Yu, G., Wang, L.-G., Han, Y., and He, Q.-Y. (2012). clusterProfiler: an R package for comparing biological themes among gene clusters. Omics 16, 284–287.

Zheng, D., Soh, B.-S., Yin, L., Hu, G., Chen, Q., Choi, H., Han, J., Chow, V.T.K., and Chen, J. (2017). Differentiation of Club Cells to Alveolar Epithelial Cells In Vitro. Sci Rep 7, 41661–41669.

